# The consequences of surviving infection across the metamorphic boundary: tradeoff insights from RNAseq and life history measures

**DOI:** 10.1101/792176

**Authors:** Naomi L.P. Keehnen, Lucie Kučerová, Sören Nylin, Ulrich Theopold, Christopher W. Wheat

**Affiliations:** Department of Zoology, Stockholm University, S-106 91 Stockholm, Sweden; Department of Molecular Biosciences, Wenner-Gren Institute, Stockholm University, Svante Arrheniusväg 20c, 10691 Stockholm, Sweden; Institute of Entomology, Biology Centre CAS, Branisovska 31, 370 05 Ceske Budejovice, Czech Republic

**Author notes:** Corresponding author: Naomi L. P. Keehnen Department of Zoology, Stockholm University, S-106 91 Stockholm, Sweden.

## Abstract

The broad diversity of insect life has been shaped, in part, by pathogen pressure, yet the influence of injury and infection during critical periods of development is understudied. During development, insects undergo metamorphosis, wherein the organism experiences a dramatic shift in their overall morphology, and physiology. In temperate zones, metamorphosis is often directly followed by a developmental arrest called diapause, for which the insect needs to acquire enough energy reserves before the onset of winter. We investigated the long-term effects of injury and infection using two bacteria in the butterfly *Pieris napi*, revealing that the negative consequences of bacterial infection carry across the metamorphic boundary. Initial direct effects of infection were weight loss and slower development, as well as an increased mortality at higher infection levels. The detrimental effects were stronger in the gram-positive *Micrococcus luteus* compared to gram-negative *Escherichia coli*. Transcriptome-wide differences between the two bacteria were already observed in the gene expression profile of the first 24 hours after infection. Larvae infected with *M. luteus* showed a strong suppression of all non-immunity related processes, with several types of immune responses being activated. The impact of these transcriptomic changes, a tradeoff between homeostasis and immune response, were visible in the life history data, wherein individuals infected with *M. luteus* had the highest mortality rate, along with the lowest pupal weight, developmental rate and adult weight of all the treatments. Overall, we find that the cost of infection and wounding in the final larval instar carries over the metamorphic boundary, and is expected to negatively affect their lifetime fitness.

## Introduction

Pathogens and parasites exert strong selection pressures upon hosts for their survival, which is exacerbated during costly developmental stages. Mounting an immune response is energetically costly in terms of physiology, development, and reproduction, and due to organism being limited by finite resources, often resulting in trade-offs between immune response and the other life history traits (Sheldon and Verhulst 1996; Freitak et al. 2003, Ahmed et al. 2002; Zuk & Stoehr 2002; Ardia et al. 2012). An energetically costly life-history trait among animals is insect metamorphosis, wherein the organism experiences a dramatic shift in their overall morphology, physiology, and often environment (e.g. from terrestrial larvae, to airborne butterfly; Russell & Dunn, 1996). Furthermore, for insect species living in temperate zones, metamorphosis is often directly followed by a developmental arrest called diapause, for which the insect needs to acquire enough energy reserves before the onset of winter (Hahn and Denlinger, 2007). Despite immunity, metamorphosis and diapause being essential and energetically costly life-history traits, the interconnection between all three has rarely been studied. An insects’ energy budget is carefully fine-tuned to integrate these traits into their life cycle. However, it is unknown how these critical resource-dependent phases of metamorphosis and diapause are affected by an infection, and what the costs are of reallocating resources to fighting this pathogen.

Insects occur in a wide variety of environments, where they are exposed to physical trauma resulting in wounds, frequent attack by parasites and pathogens, and sometimes both. The insect immune response is divided into humoral and cellular defense responses. Humoral defenses consist of the production of antimicrobial peptides via Toll and IMD signaling pathways, and enzymes like phenoloxidases (producing melanin) and reactive intermediates products (Lemaitre & Hoffmann, 2007). The cellular defenses of insects are haemocyte-mediated responses, such as phagocytosis and encapsulation (Strand, 2008). While an immune response can be effective, such a response shifts metabolic priorities within the host away from regular physiological processes, to focus on immunity and wound repair (Lochmiller and Deerenberg, 2000; Dolezal et al. 2019). As a result, the cost of immunity is not only the direct metabolic cost, but also the cost of resource allocation trade-offs (Zuk and Stoehr, 2002; Adamo et al. 2008, Ardia et al. 2011; Ardia et al. 2012). For example, developmental time, reproductive success, pupal weight, adult lifespan, all have been identified previously to trade-off with immune response, within a life stage (Boots & Begon, 1993; Thomas & Rudolf, 2010; Diamond & Kingsolver, 2011; Bajgar et al, 2015).

Any negative impact on life-history traits creates the possibility of long-term costs of successfully fighting off an infection. The complex life cycle of an insect could be delayed or disrupted, and ultimately negatively affect its lifetime survival and reproductive success. In order to fully understand the costs of the immune response, and its potential cost to other component of the organism’s life history, an integration between phenotypic and physiological analyses is needed across these complex life stages (Zuk & Stoehr, 2002; Cousteau & Chevillon 2000).

The immune response competes for resources with other energy consuming processes, like metamorphosis and diapause. Metamorphosis is a critical period in which energy stores established from larval feeding are allocated between fueling pupal development and supporting the needs of the adult for reproduction and survival (Boggs and Freeman, 2005, Boggs, 2009; Merkey et al. 2011). Despite a sharp decline in metabolic rate when entering metamorphosis, metamorphosis is a costly process, for example, in *D. melanogaster* pupae consumed 35% and 27% of their lipid and carbohydrate reserves (Merkey et al. 2011). Despite being described as a developmental arrest, diapausing individuals are not running slower than non-diapausing insects, they are following an alternative development pathway with its own unique metabolic demands (Kostal 2006, Hahn & Denlinger, 2010).

Insects that have not accumulated enough reserves to survive diapause either i) die during diapause or post-diapause development, ii) postpone diapause and try to produce one more generation, or iii) terminate diapause early when the energy reserves are low (Hahn & Denlinger, 2010). To our knowledge, it is currently unknown whether negative effects of infection and wounding during the critical larval stage influences metamorphosis and diapause. Furthermore, the majority of the immune eco-physiological studies in insects are done on a phenotypic level, measuring either the immune response, or life history characteristics. To our knowledge, no studies have looked the initial immune response on a molecular level, to see if gene expression patterns during this phase could explain life history measurements measured later in life. Transcriptome analysis can provide physiological insights into biological processes that are active in tissues, wherein a change in the expression pattern of a gene is an indication of molecular functions that are changing over time. In the case of infection studies, such insights could reveal indications of trade-offs in functional pathways. In sum, few studies have tried to integrate physiological insights via RNA-Seq with phenotypic measures of life history, across infection titers of different infection types.

Here, to gain insights into the long-term consequences of infection during critical phases of development, we immunologically challenged lepidopteran larvae preparing to pupate for diapause, after which we measured key life history traits, as well as looked at their initial transcriptomic profile of their immune response.

## Material and Methods

### Study organism and experimental design

The green veined white butterfly (*Pieris napi*) is a widespread generalist butterfly. It occurs throughout Europe and in the temperate zone of Asia (GBIF Secretariat, 2017). For this study, female butterflies were collected in northern Sweden (Abisko township) and Southern Sweden (Kullaberg park, Skåne) in August 2014. These females were transferred to Stockholm University where they were allowed to lay eggs on Garlic mustard (*Alliaria petiolata*). Larvae from wild-caught females were fed on *A. petiolata* leaves until pupation in a climate-controlled room (Light:Dark 12:12 hours, 17°C). Pupated offspring were placed in cold conditions (4°C) 21 days after pupation.

In order to test the effects of infection with gram-positive or gram-negative bacteria on survival, pupae from Abisko were taken out of diapause in April 2015 and placed in a climate-controlled room (L:D 23:1, 23°C). Unrelated males and females each received a unique identifier before release into the mating cage, and were fed ad lib on 20% sugar solution. Adults were observed every hour to ensure parentage. Once mated, females were placed in individual cups with *A. petiolata* for oviposition. The leaves were exchanged twice a day, until females stopped laying eggs. The offspring from four females that produced the highest number of eggs were chosen for the experiment. The eggs were kept in containers and placed in climate chambers to develop, and grown under diapausing conditions (L:D 8:16, 17°C). After reaching third instar, larvae were moved to individual cups containing *A. petiolata* and checked daily to monitor development.

Once larvae reached the second day of 5^th^ instar they were sexed and randomly divided among 8 treatment groups (SM figure 1). One treatment was injected with 10μl of sterilized phosphate-buffered saline (PBS), to act as a trauma control. To investigate the effect of different doses of bacteria; three treatment-groups were injected with the live gram-negative *E. coli* (10^4^, 10^5^, 10^6)^, another three treatment-groups were injected with the gram-positive *M. luteus* (10^4^, 10^5^, 10^6^), and the final treatment-group was left as uninjected controls. Details decribed below. Larvae were weighed to the nearest 0.1mg and afterwards anesthetized by chilling them in containers on ice for 5 minutes prior to injection. The syringe needle (Hamilton SYR 10uL 701 ASN) was sterilized by rinsing 3 times each in 2 tubes of 95% ethanol, followed by one tube of sterile H_2_O. The injection was done at an angle less then 45° behind the hind abdominal proleg, which was sterilized with a 95% ethanol swab beforehand (Hussa & Goodrich-Blair, 2012). Control individuals were weighed and anesthetized without injection. Larval survival was monitored twice daily, until all surviving individuals reached pupation.

For the RNA-seq experiment, larvae from Skåne, southern Sweden (Kullaberg; 56°18’N, 12°27’E 109), from the same stock as Lehmann et al. 2017, were taken out of diapause and reared in the identical conditions as above. The injection treatment was identical as above, the only deviation being the treatments, instead of eight, there were only three treatment groups: PBS, *E. coli* 10^6^, *M. luteus* 10^6^. Larva were sampled at 3, 6, 12, and, 24 hours after injection. Individuals were sampled by placing them in a 1.5 mL tube, and submerged into liquid nitrogen after which they were stored in −80c.

### Live bacteria

For both experiments, live *E. coli* DH5 alpha (1 OD = 8,3E+08 CFU/ml) and *M. luteus* CCM 169 (1 OD=1E+07 CFU/ML) were obtained from stock. The optical density (OD) was determined for both bacteria. On a daily basis, an inoculating loop was used to transfer a single colony from the LA plate to 3 ml LB broth. The culture was grown overnight at 37 °C with shaking at 250 rpm. A serial dilution was then performed to determine the number of colony forming units (CFUs) and optical density of the stock bacteria. The optical density of the broth was quantified in the spectrophotometer and used to dilute the samples to 10^4^, 10^5^ and 10^6^. The bacterial cultures were spun at 1500 rpm for 2 minutes, the supernatant was discarded and the resulting pellet resuspended using 1x PBS to obtain the 3 doses for each bacterium.

### Life history traits

For the life history experiment: survival, larva weight at second day of 5^th^ instar, time to develop to pupa after treatment, pupal mass 23 days after pupation, pupal mass 247 days after pupation, time to eclose after diapause, adult whole-body weight, abdomen weight and thorax weight were recorded (SM Figure 1). All pupae were exactly 224 days in the cold treatment, after which they were weighed to the nearest 0.1 mg and placed in a climate-controlled room (L:D 23:1 h photo cycle, 23°C). Pupae were checked twice a day to obtain accurate eclosion date. After eclosion the adults were put into 4°C for 1 day so that they could drop their meconium, after which they were weighed to obtain adult whole body, thorax and abdomen mass. Individuals were sexed in all life stages.

### Statistical analysis

Statistical analyses were performed in JMP 14 (SAS). For all analyses, data were checked for normality and heteroscedasticity where applicable. For each regression analysis (GLM), all variables were entered into the model, and non-significant variables were eliminated in a stepwise manner until the model contained only significant variables, or there was no change in the fit of the model (Akaike information criterion). Developmental rates, weights, and body ratios were investigated using Kruskal Wallis each pair comparisons. For weight data, previous studies have revealed a strong sex difference in butterflies, therefore all weight data was analyzed separately for each sex.

### Transcriptomic profiling of infection RNA isolation and sequencing

Total RNA was extracted from a total of 72 larvae, six per time point, per treatment. RNA was purified with the Direct-zol RNA MiniPrep (Zymo, CA, USA) as per manufacturer's instructions. Quality and quantity of the total RNA purified were determined using Experion equipment (Bio-Rad, CA, USA) and a Qubit instrument (Thermo Fisher Scientific, MA, USA) Due to technical error one individual of the PBS treatment failed, therefore this sampling point only has five replicates, which resulted in the final total of 71 individuals sequenced. Library preparation, sequencing and data processing of the RNA was performed at the National Genomics Infrastructure Sweden (NGI Stockholm) using strandspecific Illumina TruSeq RNA libraries with poly-A selection (Illumina HiSeq HO mode v4, paired-end 2×125 bp).

### Transcription-level expression analysis

BBduk v37.31 (https://sourceforge.net/projects/bbmap) was used to trim adapter sequences and filter to a base pair quality score of 20. Transcript-level expression analysis was done following protocol provided by Pertea et al. 2016. Briefly, reads were mapped using HISAT2 v2.1.0 (Kim et al. 2015) to the *Pieris napi* genome v1.1 (Hill et al. 2019). Samtools sort v1.7 was used to sort the file, after which it was transformed into a BAM file (Li 2009). Transcripts were assembled using StringTie v1.3.4 (Pertea et al.2016), and the *Pieris napi* v1.1 annotation file in GTF format. This resulted in an updated GTF annotation file for the *P. napi* genome. Transcript abundances were estimated for each sample using StringTie v1.3.4, and the merged transcript file as input. A gene-level read count matrix was generated using the prepDE.py script provided as part of the StringTie package, using an average read length of 125 https://ccb.jhu.edu/software/stringtie/dl/prepDE.py. Sample relationships were examined using PtR as part of Trinity v2.8.3 (Grabherr et al. 2011; Haas et al. 2013). For differential expression analysis, pairwise comparisons between all samples were conducted using DESeq2 at the gene level, including VST transformation (Love et al. 2014). Two type of DE analysis were performed, in the first analysis, genes were determined to be significantly differentially expressed when having an adjusted at a log fold change (FC) of 0, and a p-value of 0.001 or lower, representing a false discovery rate (FDR) of 0.1% on a p-value of 0.001. In the second analysis genes were determined to be significantly differentially expressed when having an adjusted at a log fold change (FC) of 2, and a p-value of 0.001 or lower.

### Cluster analysis

Two type of clustering of expression profiles over time were performed. First, to identify the overall transcription profile of the first 24 hours after infection and injury in a larva, a time series analysis was conducted genes that were differentially expressed (DEGs, (logFC 0; FDR < 0.001) between 3, 6, 12, and 24 hours after injection within each treatment (*PBS*, *E. coli,* or *M. luteus*).

Secondly, to exclude the genes being up/downregulated as a result of the injection of PBS, and to specifically identify the genes involved with the immune response, we compared the bacterial treatment with their PBS counterparts for each time point (SM Table 1). Subsequently, to investigate the expression dynamics related to the immune response over 24 hours after treatment, a time series expression cluster analysis was conducted by the bacterial treatment in comparison with the PBS treatment (logFC > 2 and logFC < −2; FDR < 0.001).

For both cluster analyses, the R package Mfuzz was used to perform the clustering using the Fuzzy c-means method (Futschik & Carlisle, 2005). First, the number of clusters were determined using K-means and the within cluster sum of squared error (SSE; elbow method) in each data set. Briefly, this method determines the sum of the squared distance between each member of a cluster and its cluster centroid, and at a certain number of clusters number the SSE will not significantly decrease with each new addition of a cluster, which provides the suitable number of clusters. Fuzzy c-means assigns each data-point a cluster membership score, where being closer to the cluster center means a higher score, and these scores are used to position the centroids. This results in a robust clustering, since low scoring data points have a reduced impact on the position of the cluster center, and as a result noise and outliers have less influence. After which the centroids were correlated to ensure that the clusters separated properly, with no correlation score above 0.85.

### GO enrichment

Gene set enrichment analysis (GSEA) was performed on the time series cluster analysis with the topGO v2.24.0 R package (Alexa et al. 2006). Genes were classified as belonging to a cluster when having a cluster score of >0.6, indicating that of all clusters, the gene belongs most to that particular cluster. The genes considered in the GSEA were those with existing GO annotations in the annotation of the genome assembly (Hill et al. 2019). In topGO, the nodeSize parameter was set to 5 to remove GO terms having fewer than five annotated genes, and other parameters were run on default. GSEA were performed using the parentchild algorithm, which takes the current parents’ terms into account. Furthermore, the ontology level was run for both biological processes and molecular function.

For the immunity time-series analysis, the DE genes were all novel transcripts constructed by StringTie, and therefore had no GO annotation in our *P. napi* v1.1 genome. Also, compared to the previous analysis, this analysis contained fewer genes. Therefore, instead of a traditional GSEA using GO terms, the genes were manually annotated. First, the exonic regions were extracted from the genome using GFFread (obtained from the Cufflinks suite at http://cole-trapnell-lab.github.io/cufflinks/; Trapnell et al. 2010). These exonic regions were searched against the Uniprot protein database (Suzek et al. 2007), using blastx (thresholds: single hit, bitscore >60 and *E*-value < 0.0001; Altschul et al., 1990).

## Results

To determine the effect of live bacteria on survivorship and Darwinian fitness proxies (e.g. weight gain, developmental rates), 5^th^ instar larvae were injected with either PBS as a control, the gram-negative *E. coli*, or the gram-positive *M. luteus* in a dose responsive manner.

### Overall survival and timing of death

In order to evaluate whether the injection itself had an effect, mortality in the PBS treatment was compared to mortality in the control treatment. Although not significant, the overall mortality was 15% higher for the group injected with PBS, compared to the non-injected control individuals (*X*^2^ = 3.46, N = 125, df = 1, P = 0.06). Next, bacterial treatment showed a significant effect on overall survival (GLM with groups (control, PBS, bacterial injections): X^2^ = 49.46, df = 10, P < 0.001, Figure 1a & SM figure 1). On average *M. luteus* elicited a higher mortality than *E. coli*, and mortality increased with an increasing dose of both pathogens (Treatment: X^2^ = 34.87, df = 7, P < 0.001; Figure 1a). For the individuals that died, timing of death was classified as either occurring during infection (death in the larval stage), during diapause (death as a pupae), or during eclosion. Overall, a higher dose of bacteria affected a more immediate death after infection, instead of mortality occurring at a later life-stage (Figure 1b; *X*^2^ = 56.47, N = 165, df = 12, P < 0.0001).

**Figure 1.**
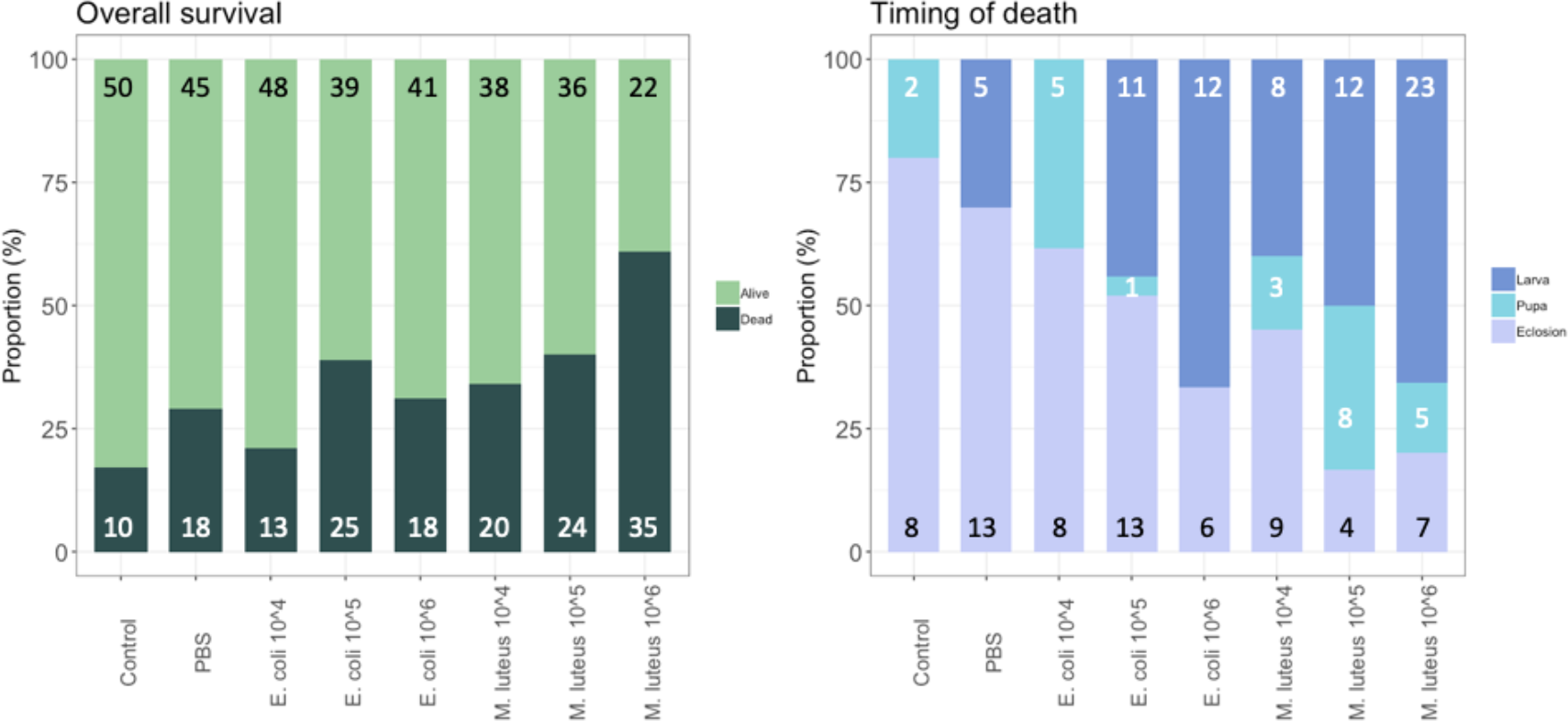
Proportion of individuals surviving and dying per treatment, with the total number per class added in the graph (left). For the individuals that died, timing of death was classified as either occurring during infection (death in the larval stage), during diapause (death as a pupae), or during eclosion (right). Graph shows the proportion of each of these groups per treatment, with the total numbers given in the graph.

### Developmental rate

For the individuals that remained alive after injections, the developmental rate, i.e. the duration of time for the larvae to pupate, was significantly different between the control and the treatments (Least Squares: F-Ratio = 8.93, df = 7, P < 0.0001, sex = n.s.). Specifically, compared to Control (M = 7.1, SD = 0.81) and PBS (M = 7.13, SD = 0.92), the infection treatments of *E. coli* 10^6^ (M = 7.95, SD = 0.84), as well as with *M. luteus*, 10^5^ (M = 8.03, SD = 1.12) and *M. luteus* 10^6^ (M = 8.45, SD = 1.37) took significantly longer to turn into pupae (Figure 2), with an average increase of up to more than a day (>18%).

**Figure 2.**
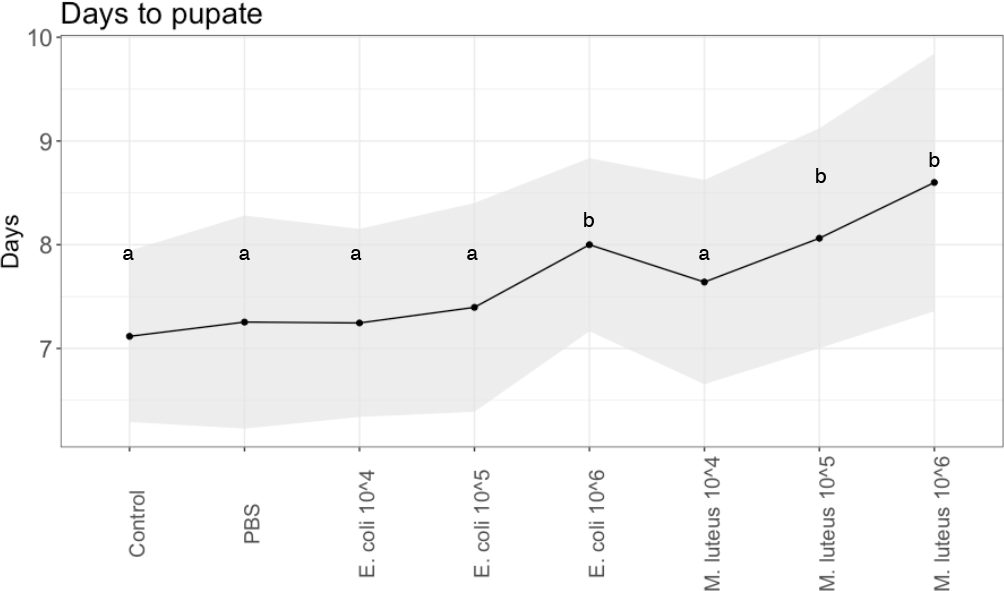
Developmental rate across injection treatments of *P. napi.* Black dots are the average per treatment, shaded areas show the standard deviations from the mean. Values not connected by the same letter are significantly different.

### Life history traits

Pupal weight was measured at two time points, first when the pupa was 23 days old (before cold treatment), and finally when the pupa was 247 days old (when they were taken out of their cold treatment to end diapause). At 23 days there was a significant difference between control individuals and most other treatments, in both males and females (Figure 3, SM Table 7). Most notably, a 13% weight loss was observed between individuals injected with PBS and individuals injected with *M. luteus* 10^6^. The pupal weight after diapause showed a similar pattern as above, with significant differences between controls and most other treatments in both sexes, as well as a 10% weight loss difference between PBS and *M. luteus* 10^6^ (Figure 3, SM Table 7).

**Figure 3.**
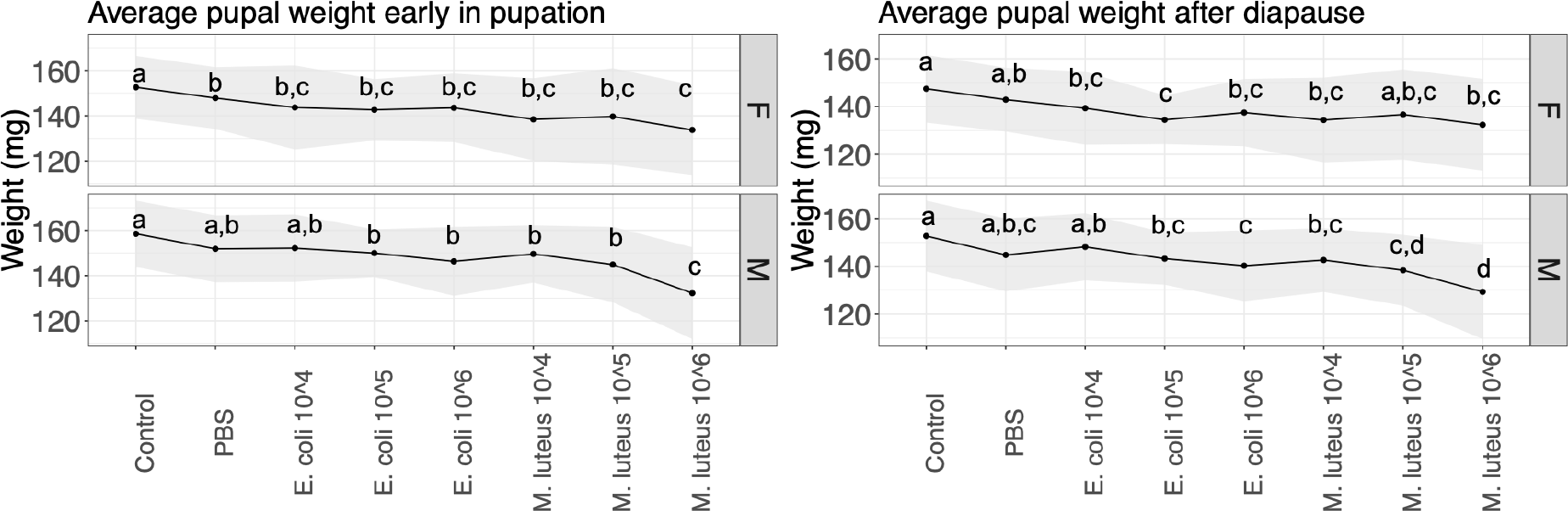
Average pupal weight over two time points per treatment. Lines show the standard deviation. Left panel shows the weight 23 days after pupation (the day they go into cold treatment), and the right panel shows the average pupa weight 247 days after pupation (stop of the cold treatment). Values not connected by the same letter are significantly different.

Adult weight after eclosion showed a significant difference between treatments in both sexes (Figure 4, Table 6). For females, all the treatments showed a significant decrease in weight compared to the controls, with the exception of M. luteus 10^5^, as well as a significant difference between PBS and *E. coli* 10^5^ (Figure 4, Table 6). For males, the differences were significant between the control and the infection treatments *E. coli* 10^5^, *E. coli* 10^6^, *M. luteus* 10^4^, and *M. luteus* 10^6^, as well as additional differences between *E. coli* 10^4^ and several other infection treatments (Figure 4, Table 6). Notably, a difference of 17% was present between PBS and *M. luteus* 10^6^ (Figure 4).

**Figure 4.**
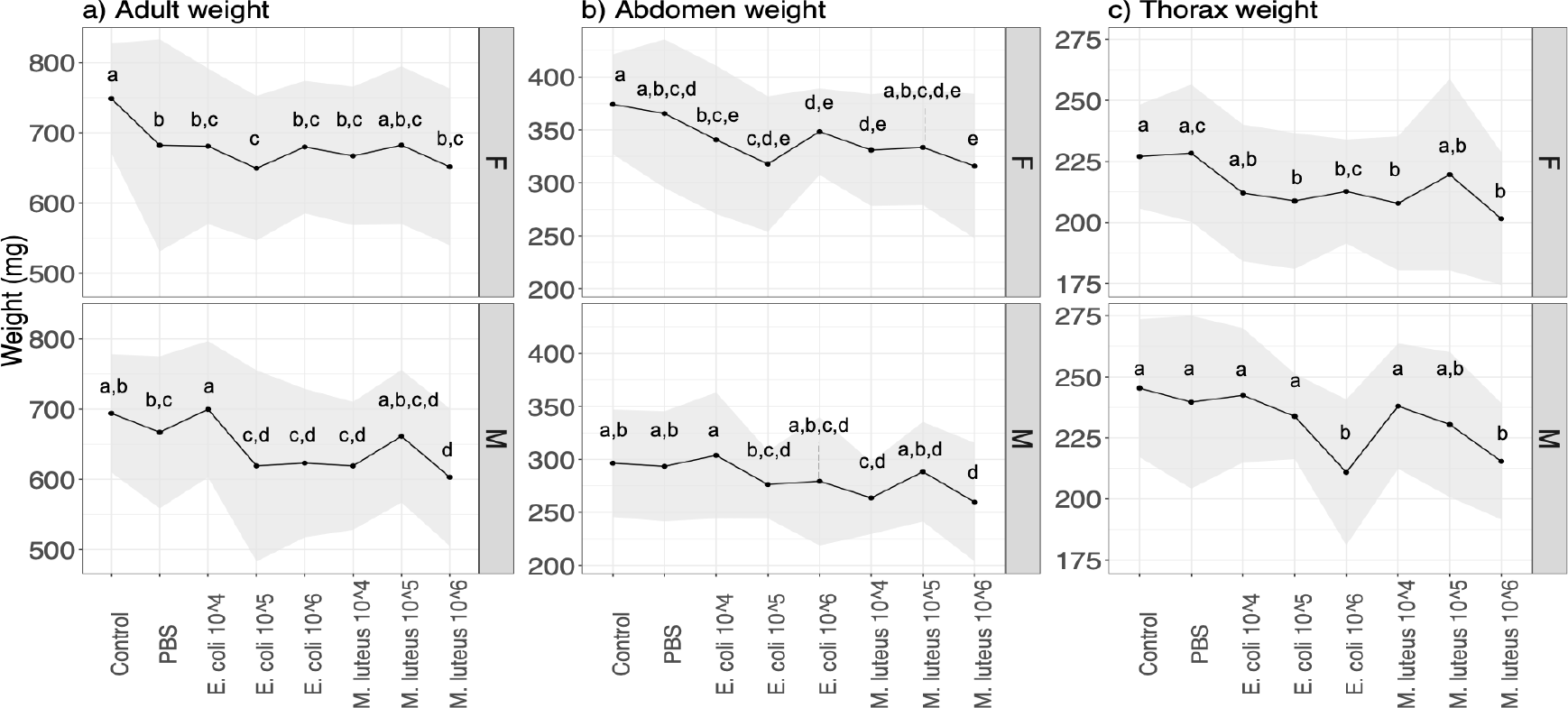
The effect of the treatment on the a) weight and b) abdomen c) thorax weight of adult butterflies. Upper panel are females (F) lower panel are males (M). The upper panels are female (F) the lower panels males (M). The shaded area denotes standard deviations. Values not connected by the same letter are significantly different.

The weight of the adult abdomen was also affected by the treatment, with female abdomen being significantly lighter in the infection treatments, compared to controls, as well as between PBS, and *E. coli* 10^5^ or *M. luteus* 10^6^ (Figure 4; SM Table 8). In males this was only true for individuals treated with *M. luteus* 10^4^, 10^6^, which differed both from controls and PBS (Figure 4, SM Table 8). Thorax weight was significantly lower in females between controls and several infection treatments, as well as between PBS and a number of infection treatments (Figure 4, SM Table 8). Males showed similar patterns (Figure 4, SM Table 8). In males treated with the highest dose of either bacteria (*E. coli 10^6^ & M. luteus* 10^6)^ had up to 12% lower thorax weight (SM Table 8).

### Transcriptome analysis of initial infection

In order to gain additional insights into the physiological responses to infection, we conducted an RNA-seq analysis across 4 time points during the first 24 hours post injections (3, 6, 12, and 24 hours), using the highest level of infection dose (10^6^). Specifically, we conducted a quantitative investigation of the transcriptome to assess the patterns emerging from the previous observations, where the highest bacterial doses of both types had the largest effects influence on immune response.

First, sample relationships were tested using a principle component analysis (PCA). This revealed that the first two PCs grouped individuals within time points per treatment, which together accounted for 49-54% of the sample variance (Figure 5). The PBS samples after 12 hours appear similar in their transcription as the 3 hours samples. The *E. coli* samples appear to get closer to their starting state after 24 hours, whereas the *M. luteus* are on a linear trajectory along PC1, with two individuals at 12 hours diverging.

**Figure 5.**
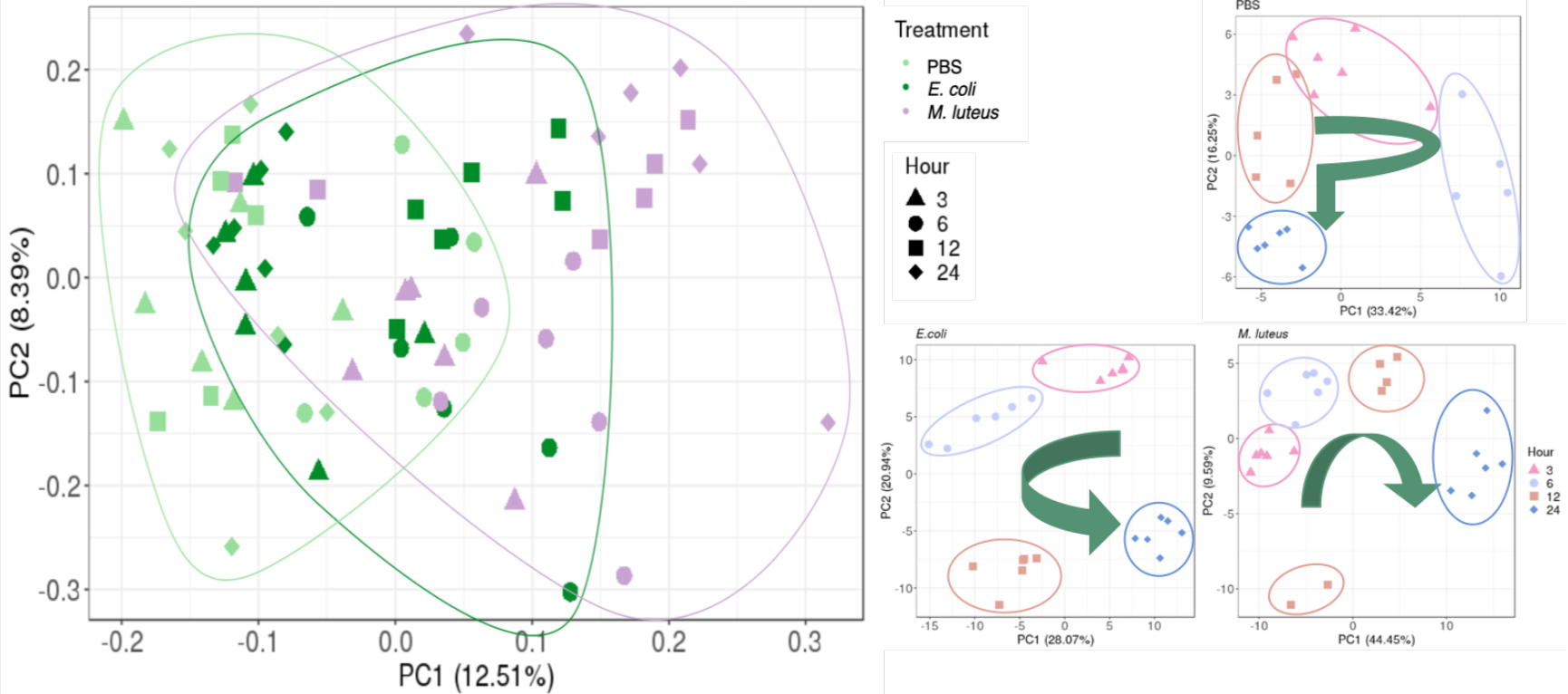
Relationship between the samples. Left graph shows the relationship of all three treatments. On the right are the comparisons of gene expression profiles when injected with PBS, *E. coli* or *M. luteus* across time. The arrows indicate the overall time progression.

### Expression dynamics over time

To investigate the expression dynamics over 24 hours after treatment, we performed a cluster analysis on the genes that were differentially expressed between any time points within each treatment. First, we determined the total number of genes differentially expressed between any time point in the experiment within each treatment (FDR < 0.001). The vast majority of DE genes were unique to each treatment, with only a small subset of genes shared by all (N=100; Figure 6).

**Figure 6.**
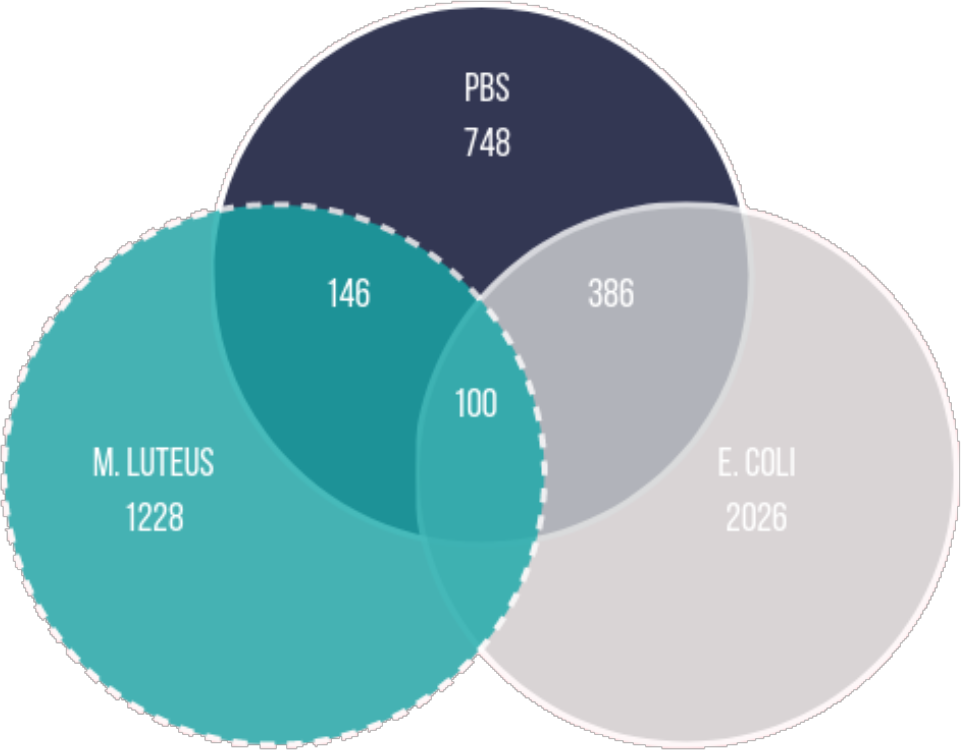
Overlap between the genes being differentially expressed in the different treatments (logFC = 0; FDR < 0.001; FC, fold change; FDR, false discovery rate)

We next grouped the differentially expressed genes in clusters using a soft clustering approach, based on the change in their expression profile over time. Cluster estimation analysis of the PBS treatment grouped the expression patterns in three clusters (Figure 7). Cluster one shows a continuous decline in expression of genes upregulated at 3 hours. GSEA revealed these genes to be involved with purine containing compound metabolism and hydrogen transport (SM Figure 2). Cluster two starts at baseline, showing higher expression at 6 hours, and a return to lower expression in the next time points. GSEA revealed genes involved with the regulation of biological process and regulation, phosphorus metabolism, and cell death (SM Figure 3). Finally, cluster 3 mirrors cluster 2, wherein it starts at baseline, but is downregulated at 6 hours, after which it gets strongly upregulated. GSEA revealed genes involved in establishment of protein localization, ncRNA metabolism and protein folding (SM Figure 4).

**Figure 7.**
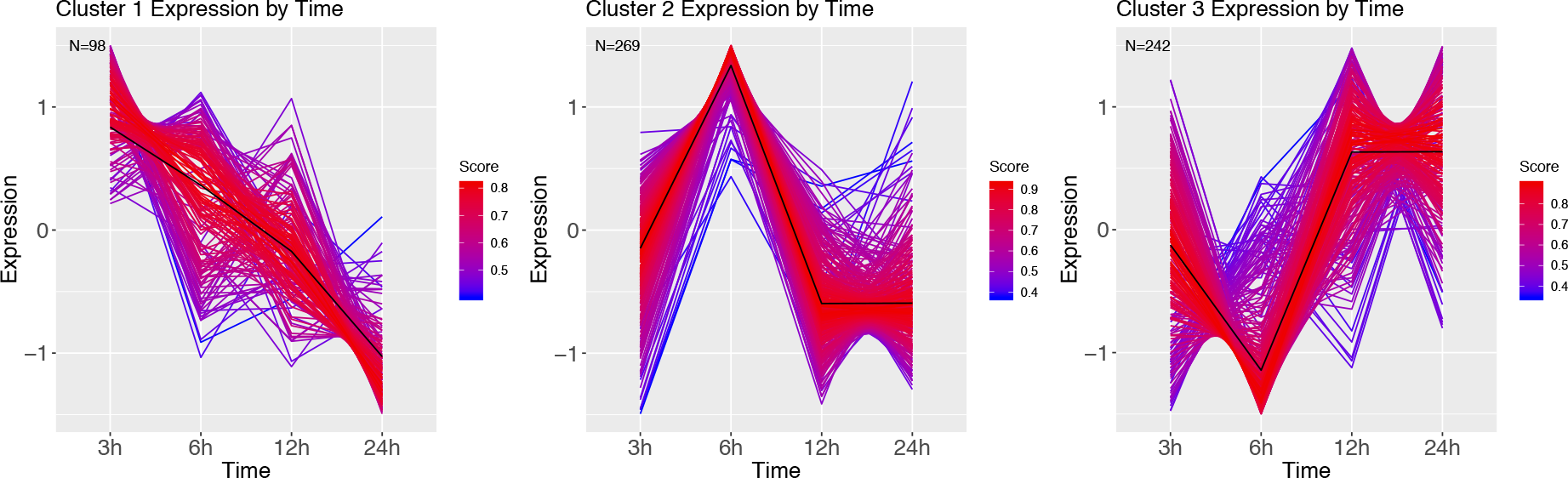
DEG clusters of larvae injected with PBS over 24 hours. Colour indicates cluster membership score, ranging from 0 (blue) to 1 (red). Numbers in each graph represent the number of genes in this cluster with a membership value higher than 0.6.

Clustering of the *E. coli* time series resulted in 6 clusters (Figure 8). Cluster 1 shows initial suppression of transcription, with upregulation starting 24 hours after infection. GSEA revealed this cluster to be related to protein alkylation and methylation (SM Figure 5). Cluster 2 mirrors 1, wherein at 6 and 12 hours after infection the genes are highly upregulated. The genes in this cluster are involved with intracellular transport, negative regulation of gene expression, and metabolism (SM Figure 6). Cluster 3 starts at 3 hours with downregulated genes, which over time get highly upregulated (at 12 hours), and at 24 hours are baseline, and contain genes involved with the regulation of cell cycle and aromatic compound biosynthesis (SM Figure 7). Cluster 4 has a strong upregulation at 6 hours, after which it becomes strongly downregulated, and contains ion transmembrane transport genes, genes involved with the regulation of biological process, cellular process (SM Figure 8.) Cluster 5 starts with upregulated genes, after which these become strongly downregulated at 12 hours, and recover to baseline 24 hours after infection. The significant GO terms associated with this cluster identified the terms: RNA modification, protein folding, biogenesis (SM Figure 9). Finally, cluster 6 goes from baseline (3hours), to downregulation (6 hours), and becomes highly upregulated the remaining two sampling points, and contains genes involved with carbohydrate metabolism, and DNA topological change (SM Figure 10).

**Figure 8.**
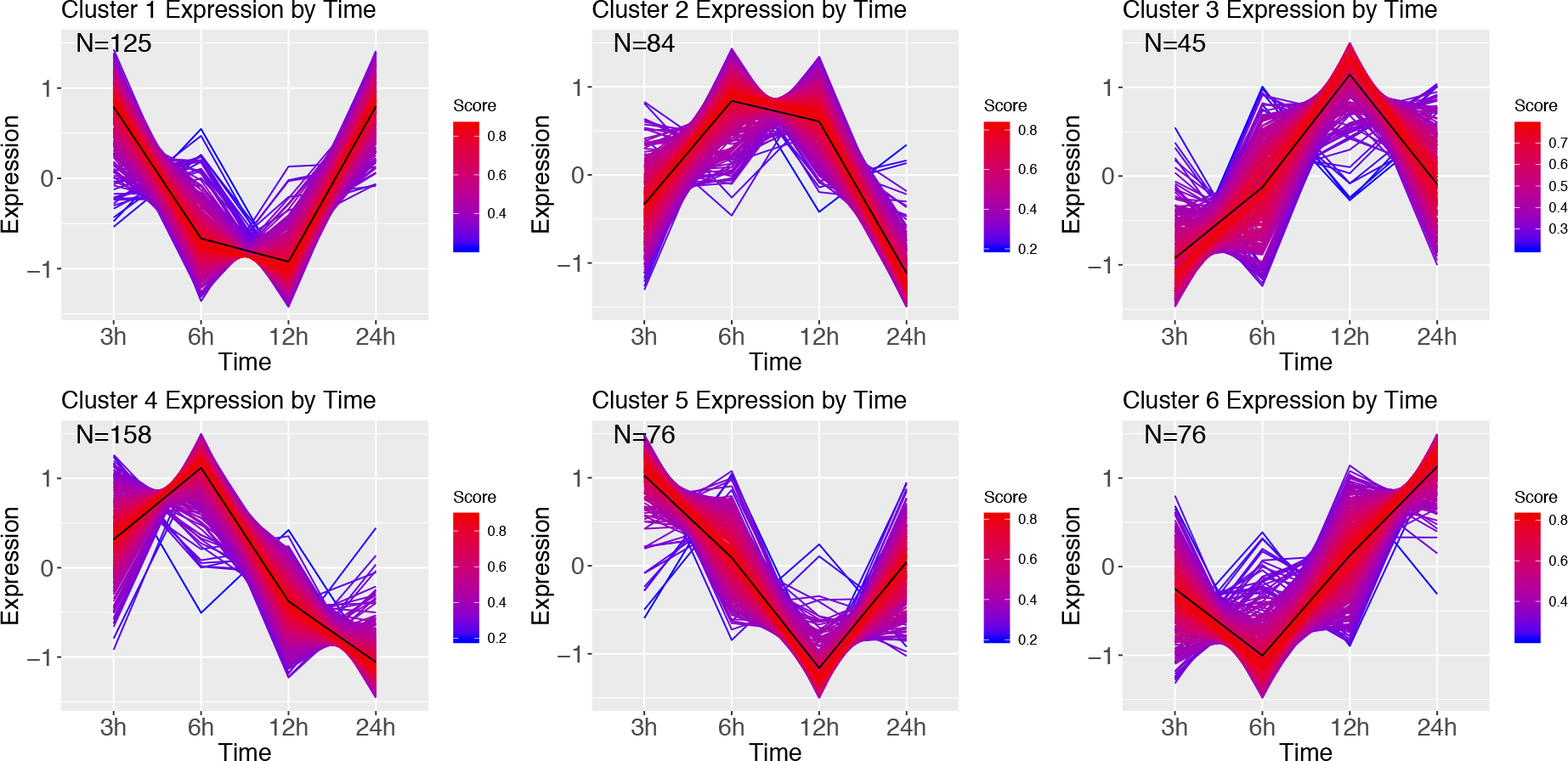
DEG clusters during *E. coli* infection over 24 hours. Colour indicates cluster membership score, ranging from 0 (blue) to 1 (red). Numbers in each graph represent the number of genes in this cluster with a membership score above 0.6.

Infection with *M. luteus* resulted in only two expression clusters over time (Figure 9). Cluster 1 show a large number of transcripts strongly upregulated during the course of infection. GSEA revealed that genes involved in the defense response and aminoglycan catabolism (SM Figure 11). Cluster 2 mirrors cluster 1, showing a large cluster of genes that are downregulated over time, and contains genes enriched for nucleoside monophosphate metabolism, metabolism and hydrogen transport (SM Figure 12).

**Figure 9.**
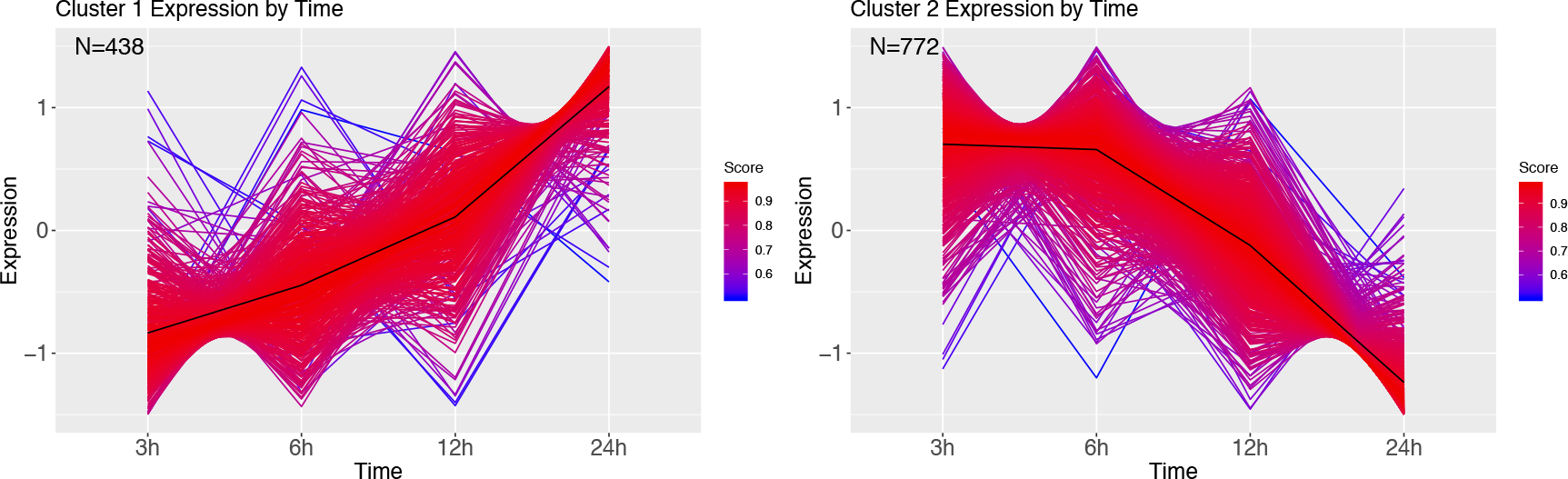
DEG clusters during *M. luteus* infection over 24 hours. Colour indicates cluster membership score, ranging from 0 (blue) to 1 (red).

### The immune response

To further investigate the expression dynamics related to the immune response over 24 hours after treatment, we did a time series expression cluster analysis by the bacterial treatment, after removing DE genes identified in the PBS time series (logFC > 2 and logFC < −2; FDR < 0.001; FC, fold change; FDR, false discovery rate). Additionally, to identify the function of the genes identified within clusters that were not annotated in our genome, all DE transcripts were searched against the uniprot protein database.

The immune response followed by an *E. coli* infection was found to have four expression clusters (Figure 10), containing 112 DE genes, of which 88 were annotated using uniprot. Cluster 1 shows downregulation of expression until 12 hours after infection, after which it is highly upregulated. Within cluster 1 there were no immune genes, but rather had an Allatostatin receptor gene and Cys-loop ligand-gated ion channel subunit-like protein (SM Table 5). Cluster 2 shows strong upregulation at 6 hours, and contained the immune genes Relish (an IMD pathway signaling gene) and Hinnavin (antimicrobial peptide; SM Table 5). Cluster 3 is downregulated at 3 hours, but shows high gene expression at 6-12 hours, after which it is back to being downregulated. The genes in this cluster with a clear immune function were antimicrobial peptides (Attacin & Moricin), and peptidoglycan recognition proteins; SM Table 5). Other genes were an Endonuclease-reverse transcriptase gene, a Chitin synthase gene, an Alkaline nuclease gene, and an actin binding protein (SM Table 5). The final cluster shows a similar pattern, but does not return to the downregulated state, and appears more baseline, and this cluster contains immune recognition genes (hemolin), modulators (serpins), as well as effector genes (antimicrobial peptides like e.g. lebocins and defensin-like peptides).

**Figure 10.**
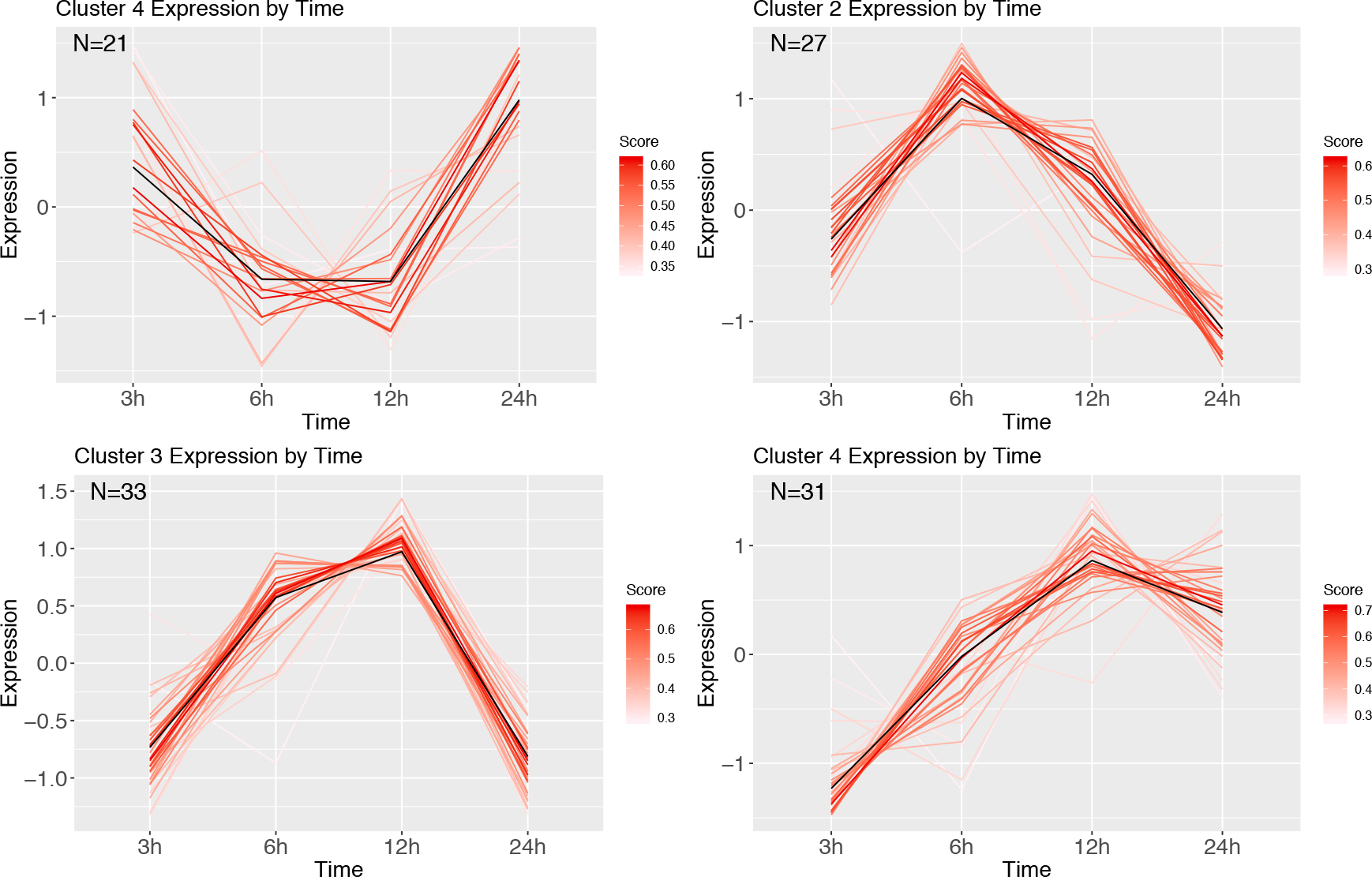
Cluster analysis over time of *E. coli* corrected with PBS. Genes were highly expressed (logFC > 2 and logFC < −2; FDR < 0.001). Each line represents a gene coloured by their membership score. Numbers in each graph represent the number of genes in this cluster.

Infection by the gram-positive *M. luteus* corrected with the PBS expression over time contained 607 DE genes, of which 463 were annotated with uniport, and were divided into three clusters (Figure 11). Cluster 1 reveals highly upregulated genes after 3 hours, which over time get strongly downregulated. Genes in this cluster were functionally diverse, but all appeared to be involved with homeostasis and metabolism. Cluster 2 contained genes that were upregulated over time and contained many immune genes. Cluster 3 starts downregulated, after which it is strongly upregulated at 6 hours after infection, and returns baseline/downregulated afterwards, and annotation revealed neutral lipase genes and sugar transporters. (need lists and tables and a bit more details here).

**Figure 11.**
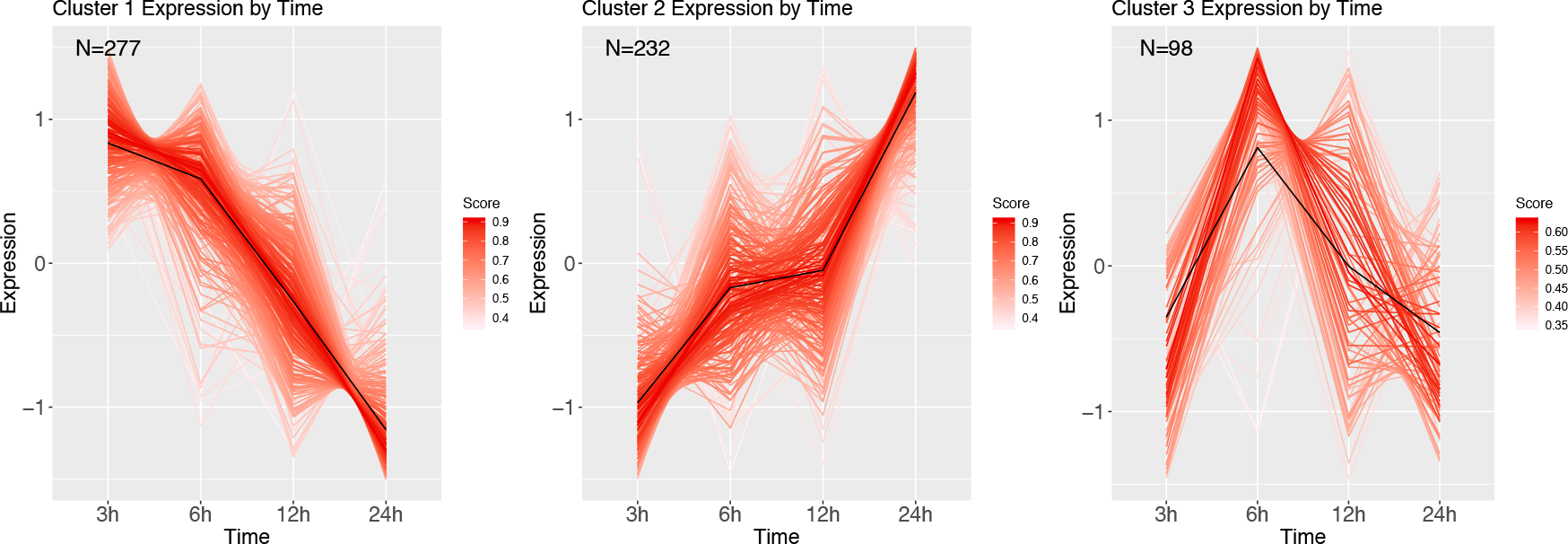
Cluster analysis over time of *M. luteus* corrected with PBS. Genes were highly expressed (logFC > 2 and logFC < −2; FDR < 0.001). Each line represents a gene coloured by their membership score. Numbers in each graph represent the number of genes in this cluster.

## Discussion

Here, we found that the negative consequences of bacterial infection carry across the metamorphic boundary in the green veined white butterfly. The type of bacteria also mattered, as the detrimental effects on life history traits were stronger in *M. luteus* compared to *E. coli*. This difference between the two bacteria was already observed in the gene expression profile during the first 24 hours after infection. Larvae infected with *M. luteus* showed a strong suppression of all non-immunity related processes, with the immune system genes being strongly upregulated. Results of this type of “overpowering” of the organism’s homeostasis was visible also in the life history data, wherein individuals infected with *M. luteus* had the highest mortality rate, along with the lowest pupal weight, developmental rate and adult weight of all the treatments.

### Overall survival

Mortality increased with an increasing dose of both pathogens and as expected, a higher dose of bacteria caused a more immediate death after infection, instead of mortality occurring at a later life-stage. Furthermore, mortality was higher with *M. luteus* than *E. coli*. These observations suggest that at lower-level infections the larvae allocate their reserves to the immune response, however, due to this reallocation, their energy reserves were significantly reduced. As a consequence of infection, mortality happening in the pupal stage suggests that these individuals died due to their inability to compensate for this loss of resources. Of the individuals that died at later stages, the majority died during eclosion, suggesting that the strain of metamorphosis was too intense, as the metamorphosis from a pupa to an adult is energetically costly. As an example, a *Manduca sexta* pupa requires 5.4 kJ of energy to complete metamorphosis, which is ~64% of the total energy available energy in lipid stores as a final instar larva that is about to pupate (Hayes et al. 1992; Odell 1998). Lipid stores are also the main fuel source used during *Drosophila melanogaster* metamorphosis, using around 35% of their total lipid store just to initiate metamorphosis (Merkey et al. 2011).

Pupal diapause brings an additional energetic demand, as the preparation for diapause, and the increased lifespan due to the delayed development, depend on the energy reserves sequestered prior to the entry into diapause (Hahn and Denlinger, 2007). In *P. napi*, lipids are the main fuel source during diapause, and lipids stores show a 74% difference (in molar percentage) going from a one-day old pupa to adult (Lehmann et al. 2016). Overall, our data showed that even if the energetic costs of an infection are met, and energetically costly metamorphosis can be completed, there are long term costs from infection, resulting in mortality due to the inability to meet the added costs of diapause.

### Hormesis at lower dose *E. coli* infection

Hormesis refers to an increase in organism performance after low level exposure to agents that are commonly harmful or toxic at higher levels of exposure (Forbes 2001). Surprisingly, during our experiment hormesis was observed, as larvae treated with a lower dose of *E. coli* showed an increased performance. Specifically, they showed the same survival rate as the control group, and had a lower mortality compared to larvae injected with PBS. Wounding causes tissue damage and the release of danger signals, and, although wounding and infection are intertwined, both show distinct signatures of gene expression (Lazzaro & Rolff, 2011; Johnston & Rolff, 2013). Our results suggest that a low-level *E. coli* infection after wounding increases survival, possibly by dual activating both wound healing and the immune response. The combination of wounding and low-level infection is likely similar to what evolutionary pressures would have responded to, since wounding and wound contamination by environmental microbes is commonplace in nature (Kamimura, 2007; Lazzaro & Rolff, 2011).

### Long term effects of larval stage infection

A slower developmental rate as a result of infection has been well documented in previous studies on insects. Interestingly, for *P. napi*, the effects of infection on the developmental rate appeared to be dependent on the bacterial type and dose. For *E. coli*, the developmental rate was significantly longer only at the highest dose, suggesting that for these gram-negative bacteria, the larvae can compensate for lower level infections and still prioritize development. However, after infection with *M. luteus* this compensation is not performed, and subsequently their developmental rate was lower regardless of bacterial dose. *E. coli* occurs in diverse forms in nature, ranging from commensal strains to those pathogenic on human or animal hosts (van Elsas et al. 2011), and further research could identify whether perhaps these bacteria, or a bacterium closely related, is common to our butterfly. Additionally, it would be interesting to study the effects of other, more ecologically relevant gram-negative and gram-positive bacteria, to see if this difference in consequences of infection between *M. luteus* and *E. coli* is a more general pattern, wherein *P. napi* perhaps have higher tolerance to gram-negative bacteria.

After metamorphosis, several negative effects of infections remained measurable. There was a significant effect of treatment on the weight of the pupae, as well as the adult butterflies. The highest dose of *E. coli*, and all dose of *M. luteus*, had significantly lower weight than the uninjected controls. However, most of the bacterial treated individuals were not significantly lower in weight compared to the PBS injected individuals. Only males that had received the highest dose of *M. luteus* had a significantly lower weight than PBS injected males. This suggests that the injury of the injection itself does not have a lasting effect, however the combination of injury and highest dose of bacterial treatment does. A classic tradeoff during a female adult butterfly life exists between flight performance and reproduction, and as a result the largest portion of an adult butterfly consists of reproductive reserves, stored in the abdomen, and flight muscle in the thorax (Boggs 1981, Wickman and Karlsson 1989 Karlsson, 1994). The abdomen of an female adult butterfly consists mostly of the reproductive organs, fat body and hemolymph, and an increase of these reserves are paralleled by a similar increase in reproductive effort for females, i.e. a larger abdomen has higher reproductive success (Wickman and Karlsson 1989). Many studies have found trade-offs between reproduction and the immune system among insects (Boots & Begon, 1993; Thomas & Rudolf, 2010; Diamond & Kingsolver, 2011). In addition to the trade-off in females, males face a similar trade-off, with thorax weight showed sensitivity to the infection treatments, potentially negatively influencing their flight capacity and/or mating resources. In sum, our data showed that infected individuals of both sexes had smaller abdomens and thoraxes. Overall, we find that the cost of infection and wounding in the final larval instar carries over the metamorphic boundary, with adults being smaller, and most likely this would affect both their flight performance as well as their reproductive output.

### Expression dynamics over time

For the gene expression analysis, larvae were injected with either PBS, or the highest dose (10^6^) of either *E. coli o*r *M. luteus*, after which sampling took place at 3, 6, 12 or 24 hours after treatment. When looking across the different gene expression profiles over time, only the larva infected with *M. luteus* show a strong signal of reallocating resources to the immune system, with a strong upregulation of genes involved with the immune system and a strong downregulation of genes involved with metabolism and organismal homeostasis. These transcriptome level observations reflect the phenotypic data, which showed a higher mortality and stronger long-term effects after exposure to *M. luteus*. Additionally, the sample relationships (Figure 5) of *M. luteus* reveals that at 12 hours after infection two individuals diverged from the others, most likely these individuals were on a trajectory to death.

The profiles of the larvae challenged by PBS, and larvae infected with *E. coli* showed more dynamic patterns, as observed in the range of expression profiles identified by the cluster analysis, had more overlapping DE genes (Figure 5), and the relationship between the samples showed to be more similar to each other than to *M. luteus* (Figure 5). Wounding and pathogen infection both activate the immune system of a host, but do so by different elicitors. A sterile wound generates exclusively danger signals, which then start the immune response. Danger signals are also present when a host is challenged by a pathogen, however, they also elicit microbe-associated molecular patterns signals (MAMPs; Lazzaro & Rolff, 2001). One possible explanation for their similarity between PBS and *E. coli* expression dynamics could be that wounding is never fully sterile, and therefore the PBS treated animals could have had some low-level infection due to this treatment, and therefore, have some MAMPs signals activating a low-level immune response. It could also be that for *P. napi* the immune challenge of *E. coli* elicits a lower immune response compared to *M. luteus*.

### Metabolism and immunity

The transcriptome of both *E. coli* and PBS showed multiple metabolic processes being up- and downregulated. However, the metabolic processes identified are known to be involved with several traits, making it challenging to interpret their role in the immune reponse of *P. napi*. Two type of metabolism we identified are potential interesting candidates for their involvement in the immunometabolism. First, during *E. coli* infection, ATP synthesis and carbohydrate metabolism both showed strong upregulation over time. Mounting an immune response is an energy-consuming process, and immune challenged individuals undergo a metabolic switch to enable the rapid production of ATP and new biomolecules via glucose and carbohydrate metabolism (Bajgar et al. 2015; Yang et al. 2017; Dolezal et al. 2019). Secondly, in the PBS treated animals, purine containing compound metabolism showed strong downregulation. High levels of purine and pyrimidine metabolites are found during the prepupal period of *Drosophila* (An et al. 2017). Perhaps this decrease in purine metabolic expression is a switch to reallocate energy previously allotted to prepupation to wound healing. However, further research is needed to confirm this hypothesis. Overall, many metabolic processes appear to be switched on and off after infection, and provide interesting candidates for future research into the immunometabolism of *P. napi*.

### The immune response

To further investigate the expression dynamics related to the immune response over 24 hours after treatment, we did a time series expression cluster analysis for each bacterial treatment, looking at the DE genes at that time point between the bacterial treatment, while accounting for wounding (via comparisons to the PBS treatment). This allowed us to investigate the physiological response uniquely attributed to bacterial exposure. The DE genes identified could be divided into the four general broad functional categories: pathogen-recognition genes (e.g., PGRPs), modulators (e.g., serpins and serine proteases), the genes of the signal transduction pathways (Toll, IMD), and effector genes encoding products that directly interact with microbes (e.g., antimicrobial peptides AMPs), or defence enzymes (pro-phenoloxidases). Activation of the insect immune system begins with the recognition of non-self through the activation of pattern recognition receptors (PRRs), encoded by recognition genes. After recognition of a pathogen, a sequence of modulation and signaling events is initiated. The Toll (Gram-positive) and IMD (Gram-negative) pathways are directly involved with the production of AMPs (Lemaitre & Hoffmann, 2007).

When comparing the genes involved with immune response against *E. coli* to those of *M. luteus*, several similarities were identified. Hemolin, a recognition gene involved with cellular immune responses, was upregulated regardless of bacterial type. Furthermore, both bacterial treatments show upregulation of x-tox proteins. X-tox genes encode immune-related proteins with imperfectly conserved tandem repeats of defensin-like motifs. In moths, however, they have lost their antimicrobial activity, suggesting they may have some other, yet unknown, function within the systemic immune response (Girard et al. 2008; Destoumieux-Garzón et al. 2009; Mikonranta et al. 2017). Additionally, both bacterial treatments upregulated several different types of antimicrobial peptides (AMPs), suggesting that to fight off the bacteria the larvae deploys a cocktail of different AMPs, that might functionally interact and synergistically attack the bacteria. As interactions between AMP’s can be achieved either via synergism, potentiation (one AMP enabling or enhancing the activity of others), or functional diversification, i.e. combinatorial activity increasing the spectrum of responses and thus the specificity of the innate immune response (Rahnamaeian et al. 2015). Both infections resulted in upregulation of the AMPs Lebocin, Moricin, and, Attacin. While both treatments showed an upregulation in common AMPs, there are a number of AMPs with treatment-specific expression profiles, produced after activation of either the Toll pathway (gram-positive bacteria) or IMD pathway (gram-negative bacteria). Heliomycin (a defensin) and lysozymes are only upregulated after *M. luteus* infection. The *E. coli* treatment showed upregulation of Hinnavin, a cecropin, which were previously found to be effective against *E. coli* (Hultmark et al. 1980). Additional variation between the expression profiles of the AMPs was found between the bacterial treatments. In the *M. luteus* treatment, all effector proteins are still strongly upregulated after 24 hours, whereas for *E. coli* the expression pattern differed between the AMPs. Hinnavin, and Attacin were strongly upregulated 6-12 hours after infection, whereas Lebocin was upregulated 12-24 hours after infection, perhaps indicative of these AMPs potentiation.

Despite having overall similarities in the genes used during the immune response, the treatments involving the two types of bacteria also showed significantly different overall expression patterns unique to a particular type. First, phenoloxidase genes were only upregulated during the *M. luteus* infection. Secondly, the immune response against *E. coli* could be divided into four expression clusters, which showed a certain level of dynamism (Figure 10), *M. luteus* only had three clusters, which broadly could be divided into genes either being strongly downregulated (metabolic, non-immunity genes), or strongly being upregulated (immunity genes) over the 24 hours (Figure 11). Interestingly, in addition to the immune genes, several other genes not directly linked to the immune system were strongly upregulated for *M. luteus*. For example, small heat shock proteins, sugar transporter proteins, cuticle proteins, and UDP-glucosyltransferase proteins showed increased expression over time. This appears to be in line with the activation of cellular immunity, which is dependent on a massive supply of glucose and glutamine (Dolezel et al. 2019). The first time-series analysis clearly revealed metabolic processes being up and down regulated for the PBS and *E. coli* treatments, however, none of these metabolic processes were unique to *E. coli*. This could imply that the effects of PBS and *E. coli* affect similar genes or allocation patterns, although it is still possible that the intensity of gene expression differs between the two treatments. In contrast, when comparing gene expression of PBS to *M. luteus*, metabolic genes showed an expression profile unique to *M. luteus*.

In sum, we found both long term and short-term effects of infection. Infection increased mortality, as well as multiple fitness parameters in individuals that survived the treatments. Furthermore, transcriptomic analysis revealed that larva infected by *M. luteus* activated several arms of the immune response, which could explain the difference in effects seen later on in the larval and adult stages. In addition, various metabolic processes were up and downregulated after both wounding and infection, providing interesting candidates for future studies looking into the immunometabolism and costs of the immune response.

## Acknowledgements

The authors would like to thank Pavel Dobes from Masaryk University in Brno, CZ for the *M. luteus* provided. Furthermore, the authors would like the thank Maria Celorio for the extraction of RNA. Johan Ljung and Pauline Gustafsson are thanked for their help during the experiments. The work was funded by the Knut and Alice Wallenberg Foundation (KAW2012.0058).

## Supplemental materials

**SM Figure 1.**
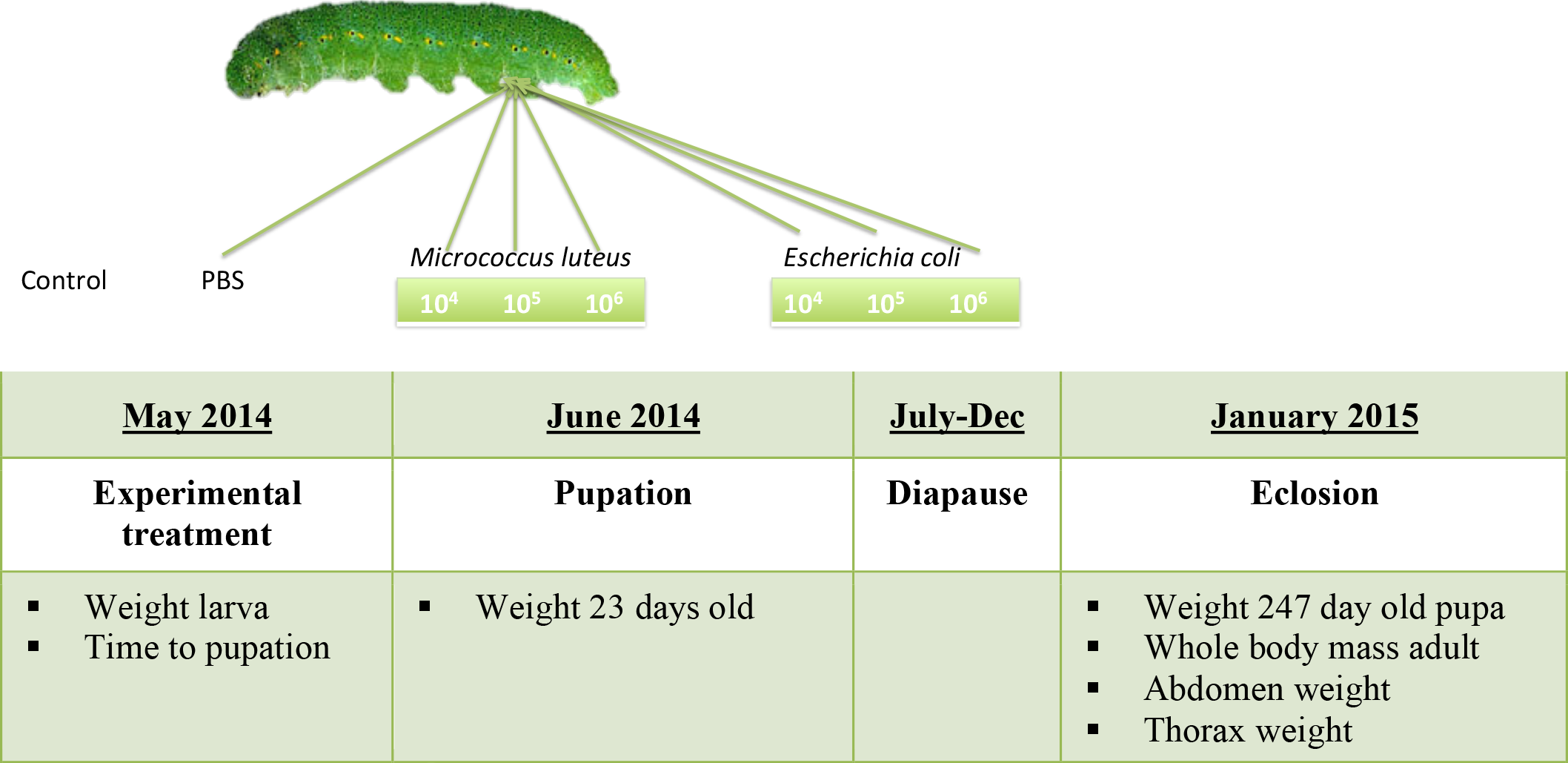
Graphical representation of the different treatments for the life history experiment.

**SM Table 1.**
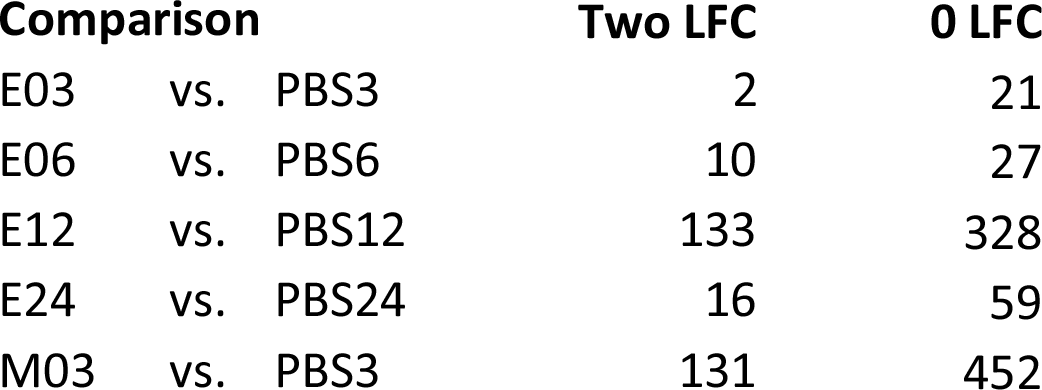
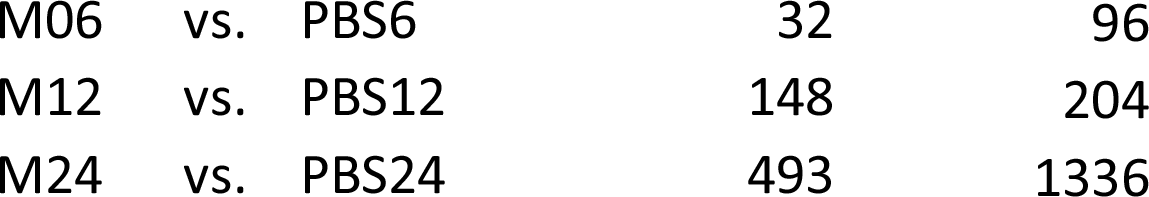
Number of significantly differential expressed genes between the two treatments at a false discovery rate < 0.001, at a log folc change of 2 and 0.

**SM Table 2.** Go term Biological processes cluster 1 *E.coli*.

| GO ID | Term | Annon. | Sign. | Exp. | classicFisher | parentchildFisher |
| --- | --- | --- | --- | --- | --- | --- |
| GO:0008213 | protein alkylation | 10 | 3 | 0,09 | 8.9e-05 | 0.0001 |
| GO:0032259 | methylation | 18 | 3 | 0,17 | 0.00058 | 0.0012 |
| GO:0043414 | macromolecule methylation | 13 | 3 | 0,12 | 0.00021 | 0.0012 |
| GO:0006396 | RNA processing | 74 | 6 | 0,7 | 5.3e-05 | 0.0021 |
| GO:0034660 | ncRNA metabolic process | 68 | 6 | 0,65 | 3.2e-05 | 0.0032 |
| GO:0044260 | cellular macromolecule metabolic process | 948 | 17 | 9 | 0.00238 | 0.0069 |
| GO:0006364 | rRNA processing | 14 | 4 | 0,13 | 6.3e-06 | 0.0081 |

**SM Table 3.** Go term Biological processes cluster 1 M. luteus.

| GO ID | Term | Annon. | Sign. | Exp. | classicFisher | parentchildFisher |
| --- | --- | --- | --- | --- | --- | --- |
| GO:0006952 | defense response | 5 | 4 | 0,14 | 2.7e-06 | 9e-05 |
| GO:0006026 | aminoglycan catabolic process | 7 | 2 | 0,19 | 0.014 | 0.0075 |
| GO:1901136 | carbohydrate derivative catabolic process | 9 | 2 | 0,25 | 0.024 | 0.0162 |

**SM Table 4.** Go term Biological processes cluster 2 M. luteus.

| GO ID | Term | Annon. | Sign. | Exp. | classicFisher | parentchildFisher |
| --- | --- | --- | --- | --- | --- | --- |
| GO:0046907 | intracellular transport | 74 | 2 | 0,35 | 0.047 | 0.016 |
| GO:0051649 | establishment of localization in cell | 86 | 2 | 0,41 | 0.061 | 0.021 |
| GO:0051641 | cellular localization | 93 | 2 | 0,44 | 0.070 | 0.025 |
| GO:0010629 | negative regulation of gene expression | 8 | 1 | 0,04 | 0.037 | 0.049 |
| GO:0010558 | negative regulation of macromolecule biosynthetic process | 8 | 1 | 0,04 | 0.037 | 0.050 |
| GO:0009890 | negative regulation of biosynthetic process | 8 | 1 | 0,04 | 0.037 | 0.050 |
| GO:2000113 | negative regulation of cellular macromolecule biosynthetic process | 8 | 1 | 0,04 | 0.037 | 0.050 |
| GO:0031327 | negative regulation of cellular biosynthetic process | 8 | 1 | 0,04 | 0.037 | 0.053 |
| GO:0008152 | metabolic process | 2218 | 14 | 10,53 | 0.051 | 0.057 |

**SM Table 5.** Results of the gene annotations done on the genes DE in *E. coli* corrected with PBS.

| gene_id | cluster | score | uniprot_name | E03 | E06 | E12 | E24 |
| --- | --- | --- | --- | --- | --- | --- | --- |
| MSTRG.24571 | 1 | 0,68 | Attacin-like antimicrobial protein | -0,85 | 0,61 | 1,09 | -0,85 |
| MSTRG.18059 | 1 | 0,66 | Endonuclease-reverse transcriptase | -0,83 | 0,70 | 1,01 | -0,89 |
| MSTRG.21526 | 1 | 0,64 | Uncharacterized protein | -0,71 | 0,58 | 1,10 | -0,97 |
| MSTRG.11624 | 1 | 0,61 | Chitin synthase | -0,90 | 0,74 | 0,98 | -0,82 |
| MSTRG.15797 | 1 | 0,60 | Antimicrobial peptide moricin | -0,94 | 0,57 | 1,11 | -0,74 |
| MSTRG.13384 | 1 | 0,59 | Alkaline nuclease | -0,63 | 0,59 | 1,09 | -1,04 |
| MSTRG.24554 | 1 | 0,57 | Attacin-like protein | -0,90 | 0,46 | 1,19 | -0,75 |
| MSTRG.3166 | 1 | 0,51 | Peptidoglycan recognition B | -1,06 | 0,62 | 1,06 | -0,62 |
| MSTRG.7993 | 1 | 0,50 | Uncharacterized protein | -0,89 | 0,89 | 0,84 | -0,84 |
| MSTRG.23753 | 1 | 0,50 | Gelsolin | -0,63 | 0,28 | 1,29 | -0,94 |
| MSTRG.18062 | 1 | 0,50 | Uncharacterized protein | -0,73 | 0,87 | 0,85 | -0,99 |
| MSTRG.3171 | 1 | 0,50 | Peptidoglycan recognition-D | -0,78 | 0,89 | 0,84 | -0,95 |
| MSTRG.10762 | 2 | 0,62 | Uncharacterized protein | -0,42 | 1,18 | 0,37 | -1,13 |
| MSTRG.22956 | 2 | 0,57 | Relish | -0,08 | 1,00 | 0,43 | -1,34 |
| MSTRG.16780 | 2 | 0,56 | BmRelish1 | 0,01 | 1,08 | 0,25 | -1,33 |
| MSTRG.12774 | 2 | 0,56 | Uncharacterized protein | 0,01 | 1,09 | 0,23 | -1,33 |
| MSTRG.15731 | 2 | 0,56 | Putative organic cation transporter | -0,16 | 1,28 | 0,04 | -1,16 |
| MSTRG.9150 | 2 | 0,55 | Uncharacterized protein | -0,24 | 0,97 | 0,56 | -1,29 |
| MSTRG.15732 | 2 | 0,55 | Putative organic cation transporter | -0,21 | 1,30 | 0,04 | -1,13 |
| MSTRG.4429 | 2 | 0,54 | Hinnavin II | -0,15 | 0,95 | 0,54 | -1,34 |
| MSTRG.11726 | 2 | 0,53 | Uncharacterized protein | 0,04 | 1,18 | 0,06 | -1,27 |
| MSTRG.12176 | 2 | 0,51 | Putative dipeptidyl-peptidase | -0,56 | 1,14 | 0,50 | -1,07 |
| MSTRG.10823 | 2 | 0,50 | Adenylate cyclase type 2 | -0,58 | 1,31 | 0,23 | -0,95 |
| MSTRG.13400 | 3 | 0,71 | Serine protease | -1,37 | -0,03 | 0,95 | 0,46 |
| MSTRG.13466 | 3 | 0,64 | Hemolin | -1,43 | 0,12 | 0,83 | 0,48 |
| MSTRG.3047 | 3 | 0,62 | Vanin-like protein 1 | -1,44 | 0,12 | 0,79 | 0,54 |
| MSTRG.23179 | 3 | 0,61 | Hemolin | -1,45 | 0,20 | 0,83 | 0,42 |
| MSTRG.3824 | 3 | 0,60 | Lebocin-like protein | -1,45 | 0,12 | 0,76 | 0,56 |
| MSTRG.12233 | 3 | 0,58 | Spod-11-tox b protein | -1,46 | 0,25 | 0,81 | 0,39 |
| MSTRG.3823 | 3 | 0,55 | Lebocin-like protein | -1,36 | -0,15 | 0,71 | 0,79 |
| MSTRG.4691 | 3 | 0,54 | Protease inhibitor-like protein | -1,19 | -0,33 | 1,17 | 0,35 |
| MSTRG.18333 | 3 | 0,54 | Serine protease | -1,20 | -0,41 | 1,02 | 0,59 |
| MSTRG.12231 | 3 | 0,54 | Antimicrobial protein 6Tox | -1,47 | 0,31 | 0,74 | 0,42 |
| MSTRG.24568 | 3 | 0,52 | Serpin-5 | -1,15 | -0,45 | 1,10 | 0,51 |
| MSTRG.7207 | 3 | 0,52 | Uncharacterized protein | -1,28 | 0,04 | 1,16 | 0,08 |
| MSTRG.3530 | 3 | 0,51 | Uncharacterized protein | -1,46 | 0,17 | 0,57 | 0,72 |
| MSTRG.14462 | 4 | 0,61 | Uncharacterized protein | 0,18 | -0,84 | -0,68 | 1,34 |
| MSTRG.4546 | 4 | 0,61 | Uncharacterized protein | 0,75 | -0,75 | -0,97 | 0,97 |
| MSTRG.20180 | 4 | 0,58 | Cys-loop ligand-gated ion channel subunit-like protein | 0,43 | -0,44 | -1,14 | 1,15 |
| MSTRG.16750 | 4 | 0,55 | Uncharacterized protein | 0,80 | -0,54 | -1,14 | 0,88 |
| MSTRG.7154 | 4 | 0,55 | Uncharacterized protein | 0,11 | -1,01 | -0,43 | 1,33 |
| MSTRG.14987 | 4 | 0,53 | Uncharacterized protein | -0,03 | -0,46 | -0,91 | 1,40 |
| MSTRG.8160 | 4 | 0,53 | Putative reverse transcriptase | 0,89 | -0,57 | -1,12 | 0,79 |
| MSTRG.4105 | 4 | 0,51 | Allatostatin receptor | -0,14 | -0,66 | -0,66 | 1,46 |

**SM Table 6.** Results of the gene annotations done on the genes DE in *M. luteus* corrected with PBS.

| gene_id | cluster | score | uniprot_name | M03 | M06 | M12 | M24 |
| --- | --- | --- | --- | --- | --- | --- | --- |
| MSTRG.13255 | 1 | 0,90 | Storage protein 1 (Fragment) | 0,96 | 0,59 | -0,26 | -1,29 |
| MSTRG.20086 | 1 | 0,90 | Phosphoserine aminotransferase | 0,97 | 0,61 | -0,34 | -1,25 |
| MSTRG.8397 | 1 | 0,90 | Cysteine synthase | 0,99 | 0,59 | -0,32 | -1,25 |
| MSTRG.16973 | 1 | 0,90 | Sorbitol dehydrogenase | 0,89 | 0,65 | -0,22 | -1,32 |
| MSTRG.20727 | 1 | 0,89 | Putative SV2-like protein 1 | 0,93 | 0,69 | -0,39 | -1,23 |
| MSTRG.7702 | 1 | 0,88 | 2-oxoglutarate dehydrogenase | 0,92 | 0,59 | -0,17 | -1,33 |
| MSTRG.8504 | 1 | 0,88 | Sodium-dependent phosphate transporter | 0,93 | 0,69 | -0,42 | -1,21 |
| MSTRG.12057 | 1 | 0,87 | Myostatin | 0,98 | 0,52 | -0,18 | -1,32 |
| MSTRG.4804 | 1 | 0,86 | Juvenile hormone esterase | 0,80 | 0,76 | -0,25 | -1,31 |
| MSTRG.1123 | 1 | 0,86 | Acyl-coa dehydrogenase | 0,93 | 0,55 | -0,13 | -1,35 |
| MSTRG.22956 | 1 | 0,86 | Nuclear factor NF-kappa-B p110 subunit | 0,82 | 0,79 | -0,35 | -1,26 |
| MSTRG.19231 | 1 | 0,85 | Phosphoglycerate kinase | 0,85 | 0,63 | -0,12 | -1,36 |
| MSTRG.17761 | 1 | 0,85 | GMP reductase | 1,07 | 0,47 | -0,31 | -1,24 |
| MSTRG.19119 | 1 | 0,85 | S-formylglutathione hydrolase | 0,96 | 0,50 | -0,12 | -1,34 |
| MSTRG.8857 | 1 | 0,85 | Antennal esterase CXE13 | 1,07 | 0,45 | -0,25 | -1,26 |
| MSTRG.21148 | 1 | 0,84 | Alcohol dehydrogenase | 1,10 | 0,45 | -0,34 | -1,21 |
| MSTRG.21152 | 1 | 0,83 | Photoreceptor dehydrogenase | 1,10 | 0,43 | -0,30 | -1,23 |
| MSTRG.8664 | 1 | 0,83 | AAEL013642-PA | 1,02 | 0,63 | -0,51 | -1,14 |
| MSTRG.25228 | 1 | 0,83 | Citrate synthase | 1,00 | 0,65 | -0,52 | -1,14 |
| MSTRG.8264 | 1 | 0,82 | Putative uncharacterized protein | 1,01 | 0,43 | -0,10 | -1,34 |
| MSTRG.1282 | 1 | 0,82 | Regucalcin | 0,81 | 0,64 | -0,06 | -1,39 |
| MSTRG.17770 | 1 | 0,82 | Glucose-6-phosphate isomerase | 0,83 | 0,61 | -0,05 | -1,39 |
| MSTRG.15184 | 1 | 0,82 | Putative uncharacterized protein | 1,11 | 0,49 | -0,46 | -1,14 |
| MSTRG.1498 | 1 | 0,81 | Neither inactivation nor afterpotential B | 1,14 | 0,42 | -0,38 | -1,18 |
| MSTRG.9500 | 1 | 0,81 | Juvenile hormone acid methyltransferase | 0,73 | 0,77 | -0,14 | -1,36 |
| MSTRG.16556 | 1 | 0,81 | Antennal esterase CXE9 | 1,15 | 0,41 | -0,39 | -1,17 |
| MSTRG.25263 | 1 | 0,80 | Citrate synthase | 1,15 | 0,43 | -0,43 | -1,15 |
| MSTRG.13740 | 1 | 0,79 | Aldo-keto reductase | 1,07 | 0,34 | -0,11 | -1,31 |
| MSTRG.19007 | 1 | 0,79 | Drongo protein isoform 2 | 0,70 | 0,88 | -0,30 | -1,28 |
| MSTRG.13246 | 1 | 0,78 | Moderately methionine rich storage protein | 0,80 | 0,60 | 0,02 | -1,42 |
| MSTRG.21721 | 1 | 0,78 | Putative nadp transhydrogenase | 0,83 | 0,55 | 0,04 | -1,42 |
| MSTRG.21720 | 1 | 0,78 | Putative nadp transhydrogenase | 0,75 | 0,66 | 0,00 | -1,41 |
| MSTRG.23484 | 1 | 0,77 | Putative igf2 mRNA binding protein | 1,19 | 0,36 | -0,44 | -1,12 |
| MSTRG.7148 | 1 | 0,77 | Putative secreted peptide 30 | 0,67 | 0,90 | -0,29 | -1,28 |
| MSTRG.16977 | 1 | 0,77 | Sorbitol dehydrogenase | 0,85 | 0,84 | -0,58 | -1,11 |
| MSTRG.9105 | 1 | 0,76 | Putative lachesin | 0,79 | 0,58 | 0,06 | -1,43 |
| MSTRG.12176 | 1 | 0,76 | Putative dipeptidyl-peptidase | 0,66 | 0,83 | -0,12 | -1,37 |
| MSTRG.5413 | 1 | 0,76 | Uricase | 1,21 | 0,28 | -0,33 | -1,16 |
| MSTRG.8211 | 1 | 0,76 | Putative fatty acid synthase | 0,80 | 0,89 | -0,55 | -1,13 |
| MSTRG.24551 | 1 | 0,76 | Putative sugar transporter | 1,16 | 0,48 | -0,59 | -1,04 |
| MSTRG.21714 | 1 | 0,75 | Putative nadp transhydrogenase | 0,89 | 0,44 | 0,09 | -1,42 |
| MSTRG.12164 | 1 | 0,75 | Putative Cyclic AMP-dependent transcription factor ATF-6 beta | 1,23 | 0,25 | -0,33 | -1,15 |
| MSTRG.11019 | 1 | 0,75 | Putative synaptic vesicle protein | 1,13 | 0,23 | -0,07 | -1,29 |
| MSTRG.8073 | 1 | 0,74 | Enoyl-CoA hydratase | 0,72 | 0,65 | 0,07 | -1,44 |
| MSTRG.20113 | 1 | 0,73 | Dipeptidyl-peptidase | 1,01 | 0,69 | -0,71 | -1,00 |
| MSTRG.15283 | 1 | 0,72 | Myo-inositol oxygenase | 1,10 | 0,58 | -0,71 | -0,98 |
| MSTRG.17341 | 1 | 0,72 | Mitochondrial aldehyde dehydrogenase | 0,90 | 0,37 | 0,14 | -1,42 |
| MSTRG.1972 | 1 | 0,72 | Alcohol dehydrogenase | 0,63 | 0,77 | 0,02 | -1,42 |
| MSTRG.15314 | 1 | 0,71 | Fructose-bisphosphate aldolase | 0,74 | 0,55 | 0,17 | -1,46 |
| MSTRG.5989 | 1 | 0,71 | Similar to CG9701-PA | 0,64 | 0,71 | 0,09 | -1,44 |
| MSTRG.17448 | 1 | 0,70 | Triosephosphate isomerase | 0,73 | 0,54 | 0,18 | -1,46 |
| MSTRG.23357 | 1 | 0,70 | Mitochondrial aldehyde dehydrogenase | 0,63 | 0,71 | 0,11 | -1,45 |
| MSTRG.21435 | 1 | 0,70 | Putative alcohol dehydrogenase | 0,65 | 0,67 | 0,13 | -1,45 |
| MSTRG.17349 | 1 | 0,70 | Mitochondrial aldehyde dehydrogenase | 0,65 | 0,67 | 0,13 | -1,45 |
| MSTRG.24584 | 1 | 0,70 | Putative sugar transporter | 1,17 | 0,49 | -0,73 | -0,93 |
| MSTRG.12014 | 1 | 0,70 | Serpin-4A | 1,05 | 0,65 | -0,78 | -0,93 |
| MSTRG.13247 | 1 | 0,69 | Moderately methionine rich storage protein | 0,71 | 0,56 | 0,20 | -1,46 |
| MSTRG.17352 | 1 | 0,69 | Aldehyde dehydrogenase (Fragment) | 0,63 | 0,68 | 0,14 | -1,45 |
| MSTRG.21355 | 1 | 0,69 | 3-hydroxyisobutyrate dehydrogenase | 0,70 | 0,58 | 0,19 | -1,46 |
| MSTRG.5990 | 1 | 0,69 | Seminal fluid protein CSSFP028 | 0,62 | 0,69 | 0,14 | -1,45 |
| MSTRG.20161 | 1 | 0,68 | Laccase 1 | 0,58 | 1,02 | -0,39 | -1,21 |
| MSTRG.9030 | 1 | 0,68 | ATP-binding cassette transporter | 0,96 | 0,76 | -0,80 | -0,93 |
| MSTRG.12427 | 1 | 0,68 | Putative sugar transporter | 0,53 | 0,90 | -0,05 | -1,38 |
| MSTRG.3035 | 1 | 0,67 | Phosphatidylethanolamine-binding protein | 0,77 | 0,43 | 0,27 | -1,47 |
| MSTRG.25711 | 1 | 0,67 | Integrin beta pat-3 | 0,91 | 0,81 | -0,80 | -0,92 |
| MSTRG.13248 | 1 | 0,67 | Moderately methionine rich storage protein | 0,89 | 0,27 | 0,27 | -1,44 |
| MSTRG.7792 | 1 | 0,66 | Endonuclease-reverse transcriptase | 1,16 | 0,51 | -0,82 | -0,85 |
| MSTRG.3034 | 1 | 0,66 | Phosphatidylethanolamine-binding protein | 0,63 | 0,63 | 0,22 | -1,47 |
| MSTRG.19202 | 1 | 0,66 | Putative argininosuccinate synthetase | 0,73 | 0,45 | 0,29 | -1,47 |
| MSTRG.21354 | 1 | 0,66 | 3-hydroxyisobutyrate dehydrogenase | 0,73 | 0,46 | 0,29 | -1,47 |
| MSTRG.10055 | 1 | 0,66 | Mo-molybdopterin cofactor sulfurase | 0,82 | 0,33 | 0,31 | -1,46 |
| MSTRG.16239 | 1 | 0,65 | Putative sugar transporter | 1,10 | 0,59 | -0,86 | -0,83 |
| MSTRG.8561 | 1 | 0,65 | Sodium-dependent phosphate transporter | 0,57 | 0,71 | 0,19 | -1,46 |
| MSTRG.13324 | 1 | 0,65 | AGAP001085-PA (Fragment) | 0,54 | 0,75 | 0,16 | -1,45 |
| MSTRG.23395 | 1 | 0,65 | CYP9G3 | 1,32 | 0,21 | -0,68 | -0,86 |
| MSTRG.21031 | 1 | 0,64 | Cellular repressor of E1A-stimulated genes | 0,50 | 0,85 | 0,08 | -1,42 |
| MSTRG.1673 | 1 | 0,64 | Follicular epithelium yolk protein subunit | 0,54 | 0,73 | 0,19 | -1,46 |
| MSTRG.22880 | 1 | 0,64 | Peritrophin type-A domain protein 3 | 1,00 | 0,72 | -0,89 | -0,83 |
| MSTRG.16240 | 1 | 0,64 | Putative sugar transporter | 1,15 | 0,53 | -0,88 | -0,80 |
| MSTRG.14709 | 1 | 0,62 | Putative lysosomal alpha-mannosidase | 0,44 | 1,02 | -0,13 | -1,33 |
| MSTRG.24743 | 1 | 0,62 | Putative Rho-associated protein kinase | 1,18 | 0,47 | -0,91 | -0,74 |
| MSTRG.25612 | 1 | 0,61 | FK506-binding protein | 0,51 | 0,70 | 0,27 | -1,48 |
| MSTRG.5445 | 1 | 0,61 | Putative venom acid phosphatase | 1,40 | 0,05 | -0,65 | -0,79 |
| MSTRG.13249 | 1 | 0,61 | Arylphorin subunit alpha | 1,41 | -0,15 | -0,32 | -0,94 |
| MSTRG.20533 | 1 | 0,61 | Putative argininosuccinate lyase | 1,42 | -0,13 | -0,41 | -0,89 |
| MSTRG.9259 | 1 | 0,60 | Putative B-cell lymphoma 3-encoded protein | 0,41 | 1,07 | -0,20 | -1,29 |
| MSTRG.8311 | 1 | 0,60 | Neuropeptide receptor A10 | 1,35 | 0,17 | -0,81 | -0,71 |
| MSTRG.12133 | 1 | 0,60 | Putative ATP-dependent RNA and DNA helicase | 1,07 | -0,09 | 0,34 | -1,32 |
| MSTRG.10384 | 2 | 0,91 | Putative topoisomerase 1-binding RING finger | -1,10 | -0,24 | 0,02 | 1,32 |
| MSTRG.12231 | 2 | 0,91 | Heli-5-tox protein | -1,05 | -0,28 | -0,02 | 1,35 |
| MSTRG.4691 | 2 | 0,91 | Protease inhibitor-like protein | -1,12 | -0,10 | -0,09 | 1,31 |
| MSTRG.19315 | 2 | 0,90 | CYP332A1 | -1,00 | -0,31 | -0,08 | 1,38 |
| MSTRG.13466 | 2 | 0,89 | Hemolin | -0,98 | -0,30 | -0,11 | 1,39 |
| MSTRG.4901 | 2 | 0,89 | VEGF27Ca | -0,99 | -0,35 | -0,03 | 1,38 |
| MSTRG.15795 | 2 | 0,88 | Moricin-like peptide C4 | -1,00 | -0,17 | -0,22 | 1,39 |
| MSTRG.24823 | 2 | 0,88 | Serine protease inhibitor 28 | -1,10 | -0,03 | -0,19 | 1,32 |
| MSTRG.23179 | 2 | 0,87 | Hemolin | -0,94 | -0,35 | -0,12 | 1,41 |
| MSTRG.24570 | 2 | 0,87 | Attacin-like antimicrobial protein | -1,08 | -0,35 | 0,13 | 1,30 |
| MSTRG.12233 | 2 | 0,87 | Antimicrobial protein 6Tox | -0,96 | -0,40 | -0,03 | 1,39 |
| MSTRG.1020 | 2 | 0,86 | Protease inhibitor 1 | -1,21 | -0,15 | 0,14 | 1,22 |
| MSTRG.10131 | 2 | 0,86 | Putative hemolymph proteinase 5 | -0,92 | -0,40 | -0,09 | 1,41 |
| MSTRG.3171 | 2 | 0,85 | Peptidoglycan recognition protein-D | -1,24 | 0,00 | 0,03 | 1,21 |
| MSTRG.24571 | 2 | 0,84 | Attacin-like antimicrobial protein | -0,91 | -0,21 | -0,31 | 1,43 |
| MSTRG.17743 | 2 | 0,84 | WAP four-disulfide core domain protein 2 | -1,26 | -0,05 | 0,12 | 1,18 |
| MSTRG.24554 | 2 | 0,84 | Attacin-like protein | -1,26 | -0,07 | 0,14 | 1,18 |
| MSTRG.6004 | 2 | 0,83 | Heat shock protein 25.4 | -0,91 | -0,17 | -0,33 | 1,42 |
| MSTRG.19735 | 2 | 0,83 | Cytochrome P450 | -1,10 | -0,39 | 0,22 | 1,26 |
| MSTRG.10902 | 2 | 0,83 | Putative cuticle protein | -1,25 | 0,06 | 0,00 | 1,19 |
| MSTRG.3249 | 2 | 0,83 | Serine protease inhibitor 27A | -1,22 | -0,22 | 0,24 | 1,19 |
| MSTRG.3823 | 2 | 0,82 | Lebocin-like protein | -1,21 | -0,24 | 0,26 | 1,19 |
| MSTRG.21819 | 2 | 0,82 | Carboxylesterase CarE-12 | -1,04 | -0,46 | 0,22 | 1,28 |
| MSTRG.10207 | 2 | 0,82 | Putative Egl nine-like protein 1 | -1,16 | -0,32 | 0,28 | 1,21 |
| MSTRG.8784 | 2 | 0,82 | Putative cuticle protein CPH43 | -0,96 | -0,52 | 0,14 | 1,34 |
| MSTRG.6262 | 2 | 0,82 | Sugar transporter | -1,13 | -0,37 | 0,28 | 1,23 |
| MSTRG.22534 | 2 | 0,81 | Small heat shock protein 27.2 | -1,08 | 0,08 | -0,32 | 1,32 |
| MSTRG.120 | 2 | 0,81 | Aldo-keto reductase | -0,93 | -0,55 | 0,14 | 1,35 |
| MSTRG.24555 | 2 | 0,81 | Attacin | -1,23 | 0,14 | -0,13 | 1,21 |
| MSTRG.6003 | 2 | 0,81 | Small heat shock protein 27.2 | -1,01 | 0,04 | -0,38 | 1,35 |
| MSTRG.14920 | 2 | 0,80 | Chemosensory protein CSP2 | -0,99 | -0,53 | 0,23 | 1,30 |
| MSTRG.24825 | 2 | 0,80 | Serine protease inhibitor 28 | -1,20 | -0,30 | 0,32 | 1,17 |
| MSTRG.3166 | 2 | 0,79 | Peptidoglycan recognition protein B | -1,04 | -0,51 | 0,29 | 1,25 |
| MSTRG.3047 | 2 | 0,79 | Vanin-like protein 1 | -1,29 | -0,12 | 0,29 | 1,12 |
| MSTRG.16505 | 2 | 0,78 | Serine protease | -1,18 | -0,36 | 0,37 | 1,16 |
| MSTRG.21817 | 2 | 0,78 | Antennal esterase CXE5 | -0,78 | -0,61 | -0,03 | 1,42 |
| MSTRG.20339 | 2 | 0,78 | Sugar transporter | -1,03 | 0,11 | -0,41 | 1,33 |
| MSTRG.10075 | 2 | 0,78 | Dynein heavy chain | -0,82 | -0,17 | -0,46 | 1,45 |
| MSTRG.11949 | 2 | 0,78 | Hemolymph proteinase 16 | -1,19 | -0,34 | 0,38 | 1,15 |
| MSTRG.3824 | 2 | 0,77 | Lebocin-like protein | -1,30 | -0,12 | 0,32 | 1,10 |
| MSTRG.7205 | 2 | 0,76 | Tetraspanin 42Ee | -1,33 | 0,12 | 0,13 | 1,09 |
| MSTRG.25490 | 2 | 0,76 | Kazal-type inhibitor | -1,30 | 0,20 | -0,03 | 1,12 |
| MSTRG.17183 | 2 | 0,76 | Chemosensory protein | -1,33 | -0,03 | 0,29 | 1,07 |
| MSTRG.19736 | 2 | 0,75 | Cytochrome CYP341A13 | -1,06 | -0,54 | 0,40 | 1,20 |
| MSTRG.121 | 2 | 0,75 | Aldo-keto reductase | -0,67 | -0,51 | -0,31 | 1,48 |
| MSTRG.9588 | 2 | 0,74 | Prophenol oxidase activating enzyme | -1,29 | 0,26 | -0,10 | 1,12 |
| MSTRG.8975 | 2 | 0,74 | Heliomicin | -0,71 | -0,73 | 0,03 | 1,40 |
| MSTRG.24425 | 2 | 0,73 | Putative alcohol dehydrogenase | -0,73 | -0,16 | -0,57 | 1,46 |
| MSTRG.22368 | 2 | 0,72 | Putative uncharacterized protein | -1,31 | -0,16 | 0,45 | 1,02 |
| MSTRG.25289 | 2 | 0,72 | Serine proteinase-like protein 1 | -1,23 | 0,33 | -0,25 | 1,15 |
| MSTRG.18744 | 2 | 0,71 | Putative cuticle protein CPH41 | -0,57 | -0,46 | -0,47 | 1,50 |
| MSTRG.25446 | 2 | 0,71 | Lysozyme II | -1,27 | -0,28 | 0,53 | 1,02 |
| MSTRG.12868 | 2 | 0,70 | Exonuclease | -1,32 | -0,17 | 0,51 | 0,98 |
| MSTRG.10823 | 2 | 0,70 | Adenylate cyclase type 2 | -0,99 | 0,26 | -0,55 | 1,28 |
| MSTRG.23724 | 2 | 0,69 | UDP-glucosyltransferase | -0,88 | -0,77 | 0,45 | 1,20 |
| MSTRG.125 | 2 | 0,69 | Aldo-keto reductase | -0,78 | -0,83 | 0,36 | 1,25 |
| MSTRG.24556 | 2 | 0,69 | Attacin-like protein | -1,29 | 0,39 | -0,17 | 1,07 |
| MSTRG.21859 | 2 | 0,68 | EP1-like protein | -1,36 | 0,35 | -0,01 | 1,01 |
| MSTRG.24552 | 2 | 0,67 | Attacin | -1,42 | 0,21 | 0,27 | 0,94 |
| MSTRG.22535 | 2 | 0,67 | Small heat shock protein 27.2 | -1,40 | 0,33 | 0,12 | 0,95 |
| MSTRG.9514 | 2 | 0,66 | EP1like2 protein | -0,54 | -0,22 | -0,71 | 1,47 |
| MSTRG.22195 | 2 | 0,65 | Putative cuticle protein CPH43 | -0,38 | -0,69 | -0,42 | 1,49 |
| MSTRG.3358 | 2 | 0,65 | UDP-glucosyltransferase | -1,01 | -0,69 | 0,67 | 1,03 |
| MSTRG.9975 | 2 | 0,64 | Alpha-esterase | -0,38 | -0,91 | -0,13 | 1,42 |
| MSTRG.21956 | 2 | 0,63 | Yellow-d | -1,10 | 0,48 | -0,51 | 1,13 |
| MSTRG.14212 | 2 | 0,62 | Serine protease inhibitor 5 | -1,15 | -0,52 | 0,77 | 0,90 |
| MSTRG.15797 | 2 | 0,62 | Antimicrobial peptide moricin | -1,20 | 0,55 | -0,39 | 1,05 |
| MSTRG.10122 | 2 | 0,61 | Putative hemolymph proteinase 5 | -0,22 | -0,85 | -0,38 | 1,45 |
| MSTRG.14358 | 3 | 0,63 | Neutral lipase | -0,71 | 1,43 | -0,05 | -0,68 |
| MSTRG.24845 | 3 | 0,62 | Zinc finger MYM-type protein 1 (Fragment) | -0,77 | 1,28 | 0,30 | -0,82 |
| MSTRG.11256 | 3 | 0,61 | Putative uncharacterized protein | -0,71 | 1,25 | 0,36 | -0,90 |
| MSTRG.15732 | 3 | 0,61 | Putative organic cation transporter | -0,77 | 1,20 | 0,44 | -0,87 |
| MSTRG.4364 | 3 | 0,61 | Ebony | -0,94 | 1,22 | 0,40 | -0,68 |
| MSTRG.6113 | 3 | 0,60 | Sugar transporter | -0,88 | 1,19 | 0,46 | -0,77 |

**SM Table 7.** Test statistics of the Kruskal Wallis each pair comparison results pupal weight per treatment. P-values that are significant are in bold.

| Comparison |  | Pupae weight before diapause |  |  |  | Pupae weight after diapause |  |  |  |
| --- | --- | --- | --- | --- | --- | --- | --- | --- | --- |
|  |  | Zfemale | p-Female | Zmale | pMale | Zfemale | p-Female | Zmale | pMale |
| E. coli 10<sup>4</sup> | Control | -2,14 | **0,032** | -1,55 | 0,119 | -2,19 | **0,028** | -1,07 | 0,281 |
| E. coli 10<sup>4</sup> | PBS | -0,54 | 0,589 | 0,31 | 0,752 | -0,65 | 0,511 | 1,26 | 0,206 |
| E. coli 10<sup>5</sup> | Control | -2,91 | **0,004** | -2,37 | **0,018** | -3,62 | **0,000** | -2,64 | **0,008** |
| E. coli 10<sup>5</sup> | E. coli 10<sup>4</sup> | -0,81 | 0,416 | -0,99 | 0,318 | -1,18 | 0,235 | -1,78 | 0,073 |
| E. coli 10<sup>5</sup> | PBS | -1,41 | 0,158 | -0,62 | 0,535 | -2,13 | **0,033** | -0,33 | 0,739 |
| E. coli 10<sup>6</sup> | Control | -2,32 | **0,020** | -2,57 | **0,010** | -2,75 | **0,006** | -2,65 | **0,008** |
| E. coli 10<sup>6</sup> | E. coli 10<sup>4</sup> | -0,18 | 0,856 | -1,57 | 0,116 | -0,31 | 0,756 | -2,23 | **0,025** |
| E. coli 10<sup>6</sup> | E. coli 10<sup>5</sup> | 0,52 | 0,602 | -0,94 | 0,343 | 0,88 | 0,375 | -0,72 | 0,467 |
| E. coli 10<sup>6</sup> | PBS | -0,75 | 0,451 | -1,24 | 0,214 | -1,15 | 0,250 | -0,87 | 0,383 |
| M. luteus 10<sup>4</sup> | Control | -2,78 | **0,005** | -2,09 | **0,036** | -2,60 | **0,009** | -2,39 | **0,016** |
| M. luteus 10<sup>4</sup> | E. coli 10<sup>4</sup> | -0,89 | 0,373 | -1,04 | 0,297 | -0,48 | 0,631 | -1,91 | 0,055 |
| M. luteus 10<sup>4</sup> | E. coli 10<sup>5</sup> | -0,30 | 0,764 | 0,06 | 0,951 | 0,58 | 0,557 | -0,09 | 0,923 |
| M. luteus 10<sup>4</sup> | E. coli 10<sup>6</sup> | -0,71 | 0,475 | 0,59 | 0,554 | -0,25 | 0,802 | 0,37 | 0,707 |
| M. luteus 10<sup>4</sup> | PBS | -1,31 | 0,188 | -0,60 | 0,542 | -1,10 | 0,268 | -0,43 | 0,663 |
| M. luteus 10<sup>5</sup> | Control | -2,15 | **0,031** | -2,82 | **0,005** | -1,81 | 0,069 | -3,07 | **0,002** |
| M. luteus 10<sup>5</sup> | E. coli 10<sup>4</sup> | -0,65 | 0,510 | -1,57 | 0,115 | -0,42 | 0,669 | -2,45 | **0,014** |
| M. luteus 10<sup>5</sup> | E. coli 10<sup>5</sup> | 0,036 | 0,971 | -1,06 | 0,287 | 0,58 | 0,560 | -1,19 | 0,233 |
| M. luteus 10<sup>5</sup> | E. coli 10<sup>6</sup> | -0,46 | 0,642 | 0,01 | 0,991 | -0,03 | 0,969 | 0,02 | 0,981 |
| M. luteus 10<sup>5</sup> | M. luteus 10<sup>4</sup> | 0,40 | 0,682 | -0,83 | 0,407 | 0,27 | 0,786 | -1,00 | 0,3173 |
| M. luteus 10<sup>5</sup> | PBS | -1,10 | 0,268 | -1,40 | 0,161 | -0,86 | 0,386 | -1,42 | 0,1534 |
| M. luteus 10<sup>6</sup> | Control | -3,19 | **0,001** | -4,14 | **<0,0001** | -2,59 | **0,009** | -3,87 | **0,0001** |
| M. luteus 10<sup>6</sup> | E. coli 10<sup>4</sup> | -1,82 | 0,068 | -3,74 | **0,000** | -0,95 | 0,338 | -3,81 | **0,0001** |
| M. luteus 10<sup>6</sup> | E. coli 10<sup>5</sup> | -1,70 | 0,089 | -3,35 | **0,001** | -0,50 | 0,613 | -2,73 | **0,0062** |
| M. luteus 10<sup>6</sup> | E. coli 10<sup>6</sup> | -1,82 | 0,068 | -2,28 | **0,022** | -0,80 | 0,423 | -1,87 | 0,0603 |
| M. luteus 10<sup>6</sup> | M. luteus 10<sup>4</sup> | -0,96 | 0,337 | -3,19 | **0,001** | -0,60 | 0,543 | -2,51 | **0,0119** |
| M. luteus 10<sup>6</sup> | M. luteus 10<sup>5</sup> | -1,10 | 0,271 | -1,94 | 0,052 | -0,62 | 0,531 | -1,09 | 0,2721 |
| M. luteus 10<sup>6</sup> | PBS | -2,25 | **0,024** | -3,48 | **0,001** | -1,77 | 0,077 | -2,64 | **0,0082** |
| PBS | Control | -1,96 | **0,050** | -1,58 | 0,113 | -1,8 | 0,058 | -1,93 | 0,0535 |

**SM Table 8.** Test statistics results of the Kruskal Wallis each pair comparison Adult body weight, Thorax Abdomen. P-values that are significant are in bold.

| Comparison |  | Whole body weight |  |  |  | Abdomenweight |  |  |  | Thorax weight |  |  |  |
| --- | --- | --- | --- | --- | --- | --- | --- | --- | --- | --- | --- | --- | --- |
| Level | - Level | Zfemale | p-Female | Zmale | pMale | Zfemale | p-Female | Zmale | pMale | Zfemale | p-Female | Zmale | pMale |
| E. coli 10<sup>4</sup> | Control | -2,29 | **0,0219** | 0,33 | 0,735 | -1,97 | **0,048** | 0,95 | 0,339 | -1,68 | 0,092 | -0,19 | 0,842 |
| E. coli 10<sup>4</sup> | PBS | -0,79 | 0,426 | 1,66 | 0,097 | -0,79 | 0,424 | 1,02 | 0,306 | -1,79 | 0,073 | 0,14 | 0,887 |
| E. coli 10<sup>5</sup> | Control | -3,43 | **0,0006** | -2,40 | **0,016** | -2,97 | **0,003** | -1,75 | 0,079 | -2,42 | **0,015** | -1,71 | 0,087 |
| E. coli 10<sup>5</sup> | E. coli 10<sup>4</sup> | -0,80 | 0,4212 | -2,74 | **0,006** | -1,29 | 0,196 | -2,76 | **0,005** | -0,28 | 0,778 | -1,71 | 0,087 |
| E. coli 10<sup>5</sup> | PBS | -2,17 | **0,0299** | -0,59 | 0,555 | -2,03 | **0,041** | -1,02 | 0,306 | -1,98 | **0,047** | -0,92 | 0,353 |
| E. coli 10<sup>6</sup> | Control | -2,65 | **0,0079** | -2,41 | **0,016** | -2,07 | **0,038** | -1,31 | 0,188 | -2,44 | **0,015** | -3,16 | **0,002** |
| E. coli 10<sup>6</sup> | E. coli 10<sup>4</sup> | 0,13 | 0,8919 | -2,90 | **0,004** | 0,26 | 0,790 | -1,79 | 0,073 | -0,08 | 0,932 | -3,51 | **0,000** |
| E. coli 10<sup>6</sup> | E. coli 10<sup>5</sup> | 1,14 | 0,2504 | -0,83 | 0,404 | 1,76 | 0,077 | -0,09 | 0,924 | 0,56 | 0,571 | -2,61 | **0,009** |
| E. coli 10<sup>6</sup> | PBS | -1,03 | 0,2988 | -1,41 | 0,156 | -0,39 | 0,690 | -0,94 | 0,343 | -1,93 | 0,053 | -2,64 | **0,008** |
| M. luteus 10<sup>4</sup> | Control | -2,98 | **0,0028** | -2,84 | **0,004** | -2,60 | **0,009** | -2,50 | **0,012** | -2,17 | **0,030** | -0,63 | 0,527 |
| M. luteus 10<sup>4</sup> | E. coli 10<sup>4</sup> | -0,44 | 0,6532 | -3,14 | **0,002** | -0,47 | 0,636 | -3,31 | **0,001** | -0,56 | 0,569 | -0,94 | 0,346 |
| M. luteus 10<sup>4</sup> | E. coli 10<sup>5</sup> | 0,43 | 0,6648 | -0,76 | 0,445 | 0,65 | 0,514 | -1,50 | 0,132 | 0 | 1,000 | 0,33 | 0,740 |
| M. luteus 10<sup>4</sup> | E. coli 10<sup>6</sup> | -0,60 | 0,5466 | 0,24 | 0,808 | -1,03 | 0,302 | -0,89 | 0,373 | -0,43 | 0,666 | 2,35 | **0,019** |
| M. luteus 10<sup>4</sup> | PBS | -1,32 | 0,1846 | -1,26 | 0,207 | -1,42 | 0,154 | -2,17 | **0,029** | -2,15 | **0,031** | -0,17 | 0,860 |
| M. luteus 10<sup>5</sup> | Control | -1,94 | 0,0523 | -1,15 | 0,250 | -2,26 | **0,023** | -0,17 | 0,860 | -0,31 | 0,756 | -1,64 | 0,099 |
| M. luteus 10<sup>5</sup> | E. coli 10<sup>4</sup> | 0,14 | 0,8872 | -1,18 | 0,235 | -0,29 | 0,770 | -1,18 | 0,234 | 0,60 | 0,548 | -1,43 | 0,150 |
| M. luteus 10<sup>5</sup> | E. coli 10<sup>5</sup> | 1,03 | 0,2992 | 1,37 | 0,169 | 0,72 | 0,470 | 1,40 | 0,160 | 0,89 | 0,372 | -0,05 | 0,959 |
| M. luteus 10<sup>5</sup> | E. coli 10<sup>6</sup> | 0,13 | 0,8917 | 1,50 | 0,131 | -0,86 | 0,390 | 1,01 | 0,312 | 0,66 | 0,506 | 1,94 | 0,052 |
| M. luteus 10<sup>5</sup> | M. luteus 10<sup>4</sup> | 0,43 | 0,6616 | 1,69 | 0,091 | 0,05 | 0,956 | 2,14 | **0,031** | 1,02 | 0,308 | -0,73 | 0,463 |
| M. luteus 10<sup>5</sup> | PBS | -0,56 | 0,5706 | 0,07 | 0,938 | -1,33 | 0,183 | 0,06 | 0,947 | -0,56 | 0,570 | -1,01 | 0,312 |
| M. luteus 10<sup>6</sup> | Control | -2,48 | **0,013** | -2,86 | **0,004** | -2,55 | **0,011** | -2,21 | **0,027** | -2,42 | **0,015** | -2,83 | **0,005** |
| M. luteus 10<sup>6</sup> | E. coli 10<sup>4</sup> | -0,67 | 0,4969 | -3,09 | **0,002** | -1,05 | 0,292 | -2,60 | **0,009** | -0,82 | 0,412 | -3,28 | **0,001** |
| M. luteus 10<sup>6</sup> | E. coli 10<sup>5</sup> | 0 | 1 | -1,57 | 0,116 | -0,12 | 0,898 | -1,78 | 0,074 | -0,56 | 0,571 | -2,41 | **0,016** |
| M. luteus 10<sup>6</sup> | E. coli 10<sup>6</sup> | -0,92 | 0,3525 | -0,85 | 0,395 | -1,65 | 0,098 | -1,22 | 0,222 | -1,06 | 0,286 | 0,03 | 0,971 |
| M. luteus 10<sup>6</sup> | M. luteus 10<sup>4</sup> | -0,58 | 0,5563 | -1,25 | 0,210 | -0,93 | 0,350 | -0,96 | 0,334 | -0,41 | 0,676 | -2,20 | **0,028** |
| M. luteus 10<sup>6</sup> | M. luteus 10<sup>5</sup> | -0,72 | 0,4688 | -1,78 | 0,074 | -0,97 | 0,329 | -1,87 | 0,060 | -1,26 | 0,208 | -1,50 | 0,132 |
| M. luteus 10<sup>6</sup> | PBS | -1,56 | 0,1172 | -1,96 | **0,050** | -2,06 | **0,039** | -2,05 | **0,040** | -2,22 | **0,026** | -2,18 | **0,029** |
| PBS | Control | -1,98 | **0,0468** | -1,04 | 0,296 | -1,37 | 0,168 | -0,15 | 0,877 | 0,16 | 0,872 | -0,52 | 0,598 |

**SM Figure 2.**
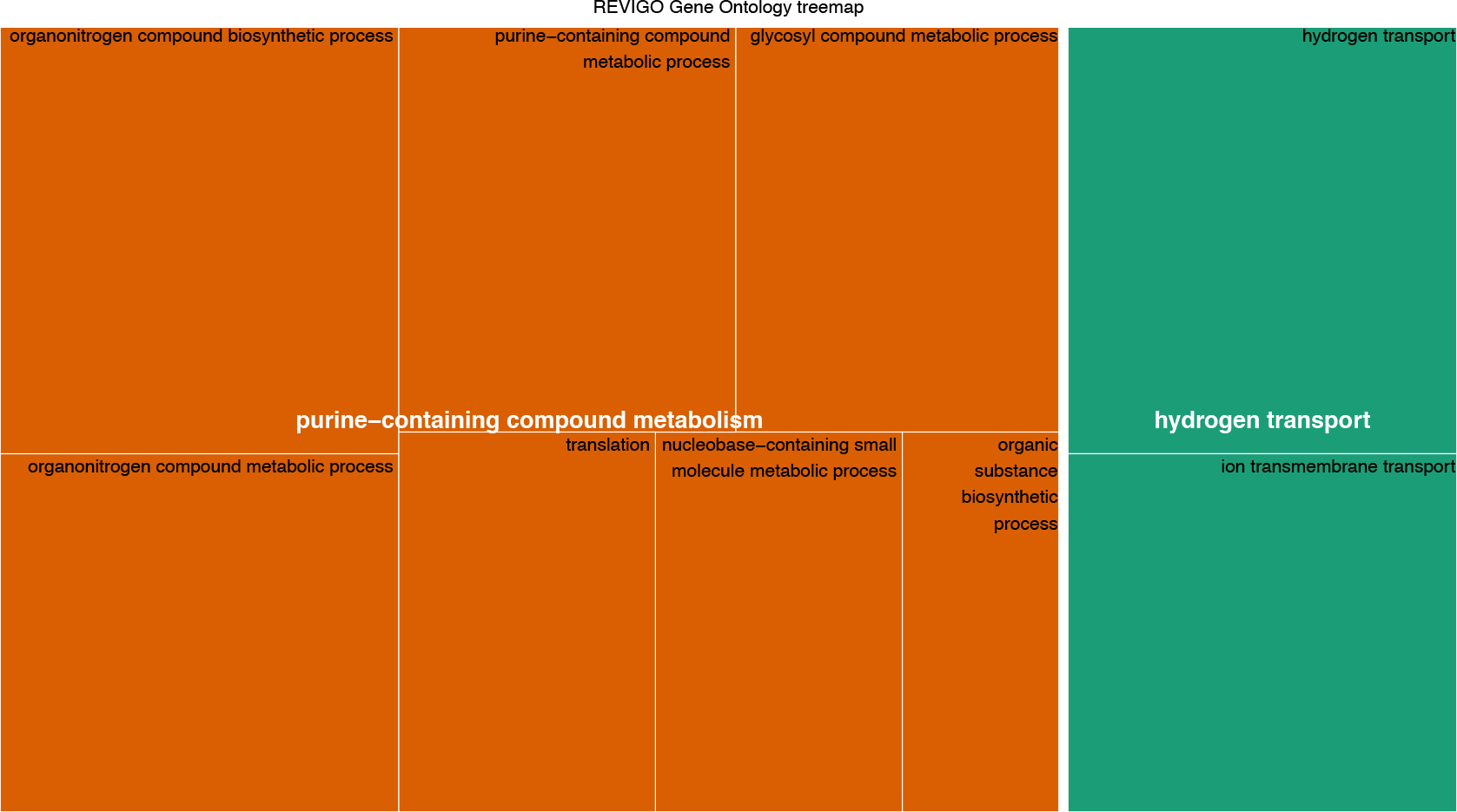
PBS cluster 1.

**SM Figure 3.**
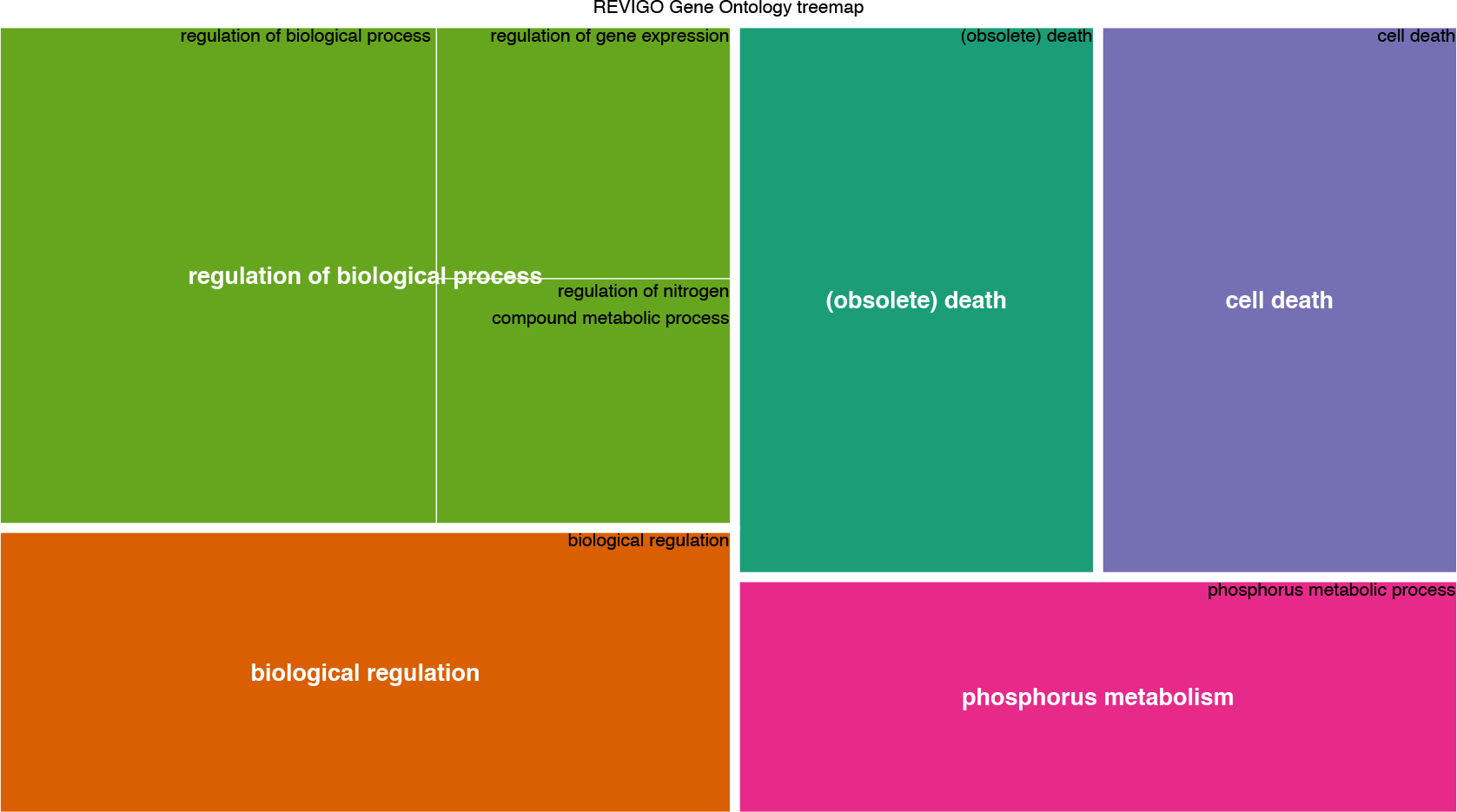
PBS cluster 2.

**SM Figure 4.**
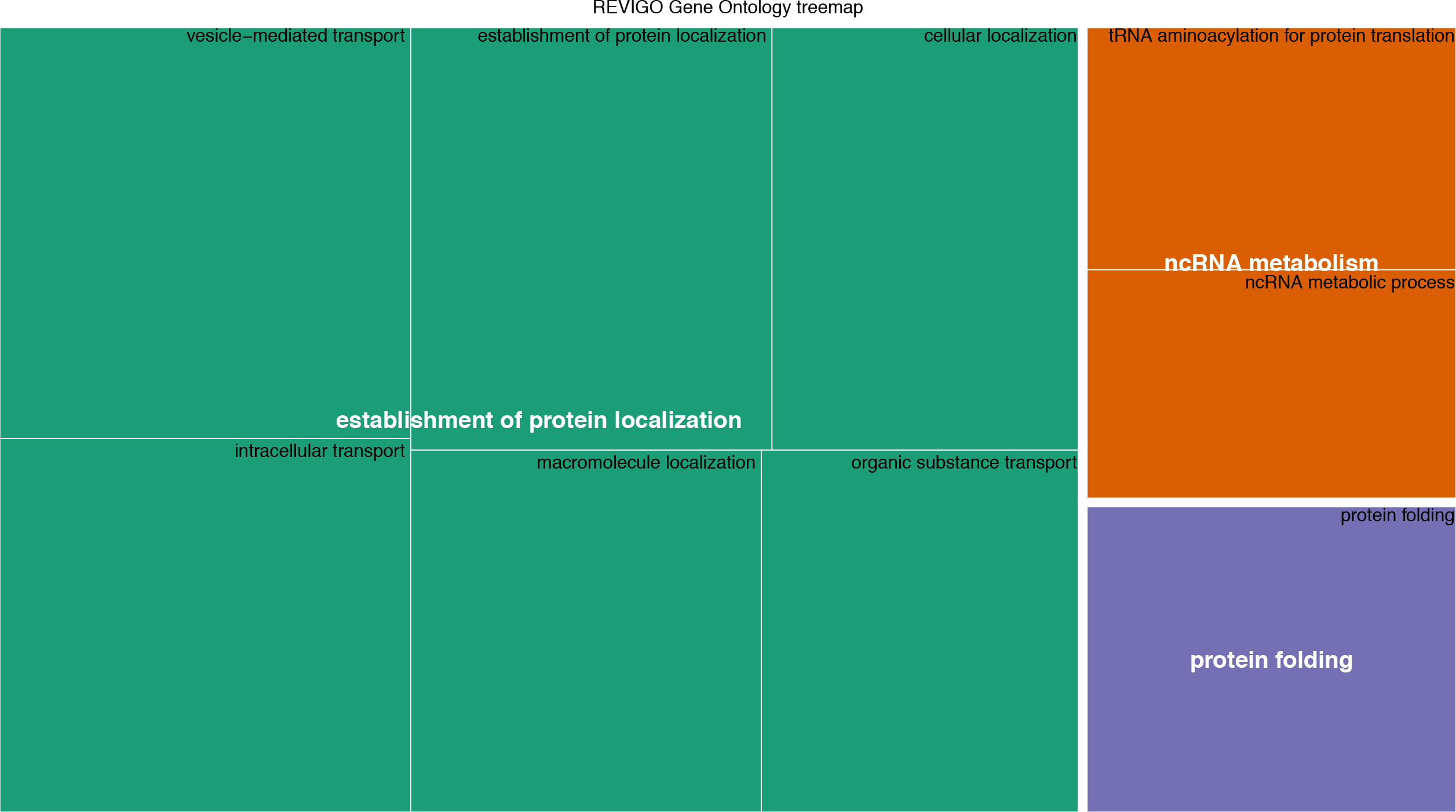
PBS cluster 3.

**SM Figure 5.**
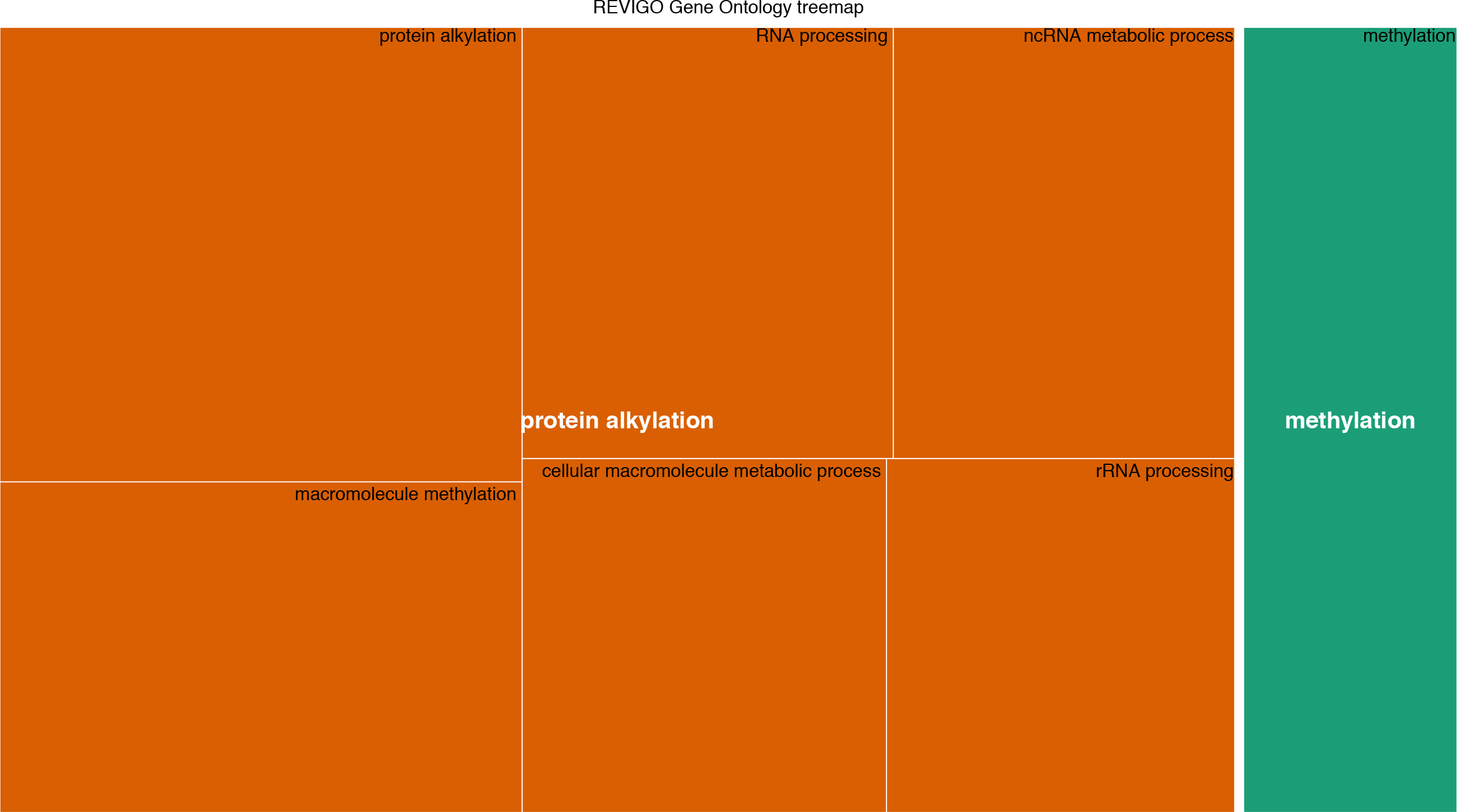
E. coli Cluster 1.

**SM Figure 6.**
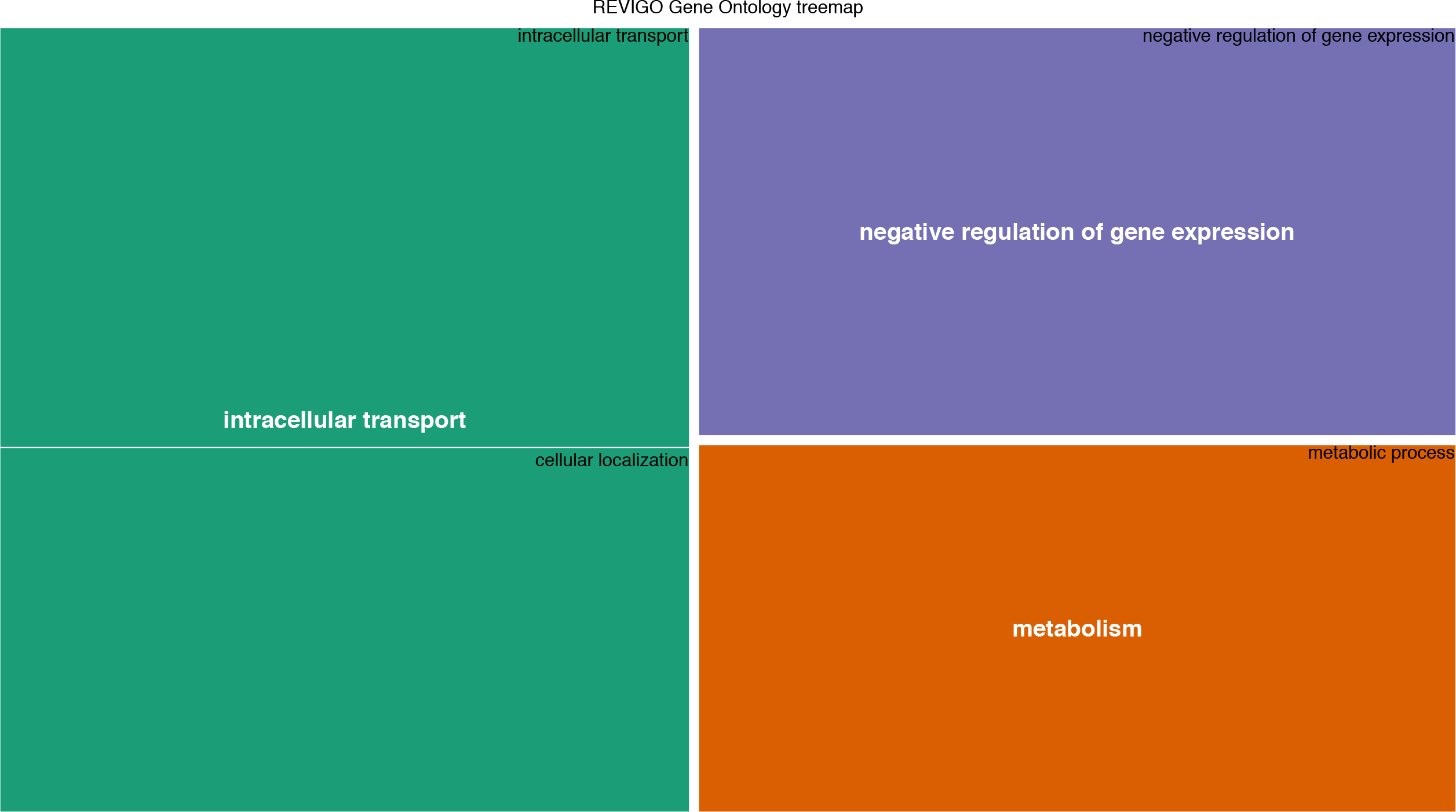
E. coli cluster 2.

**SM Figure 7.**
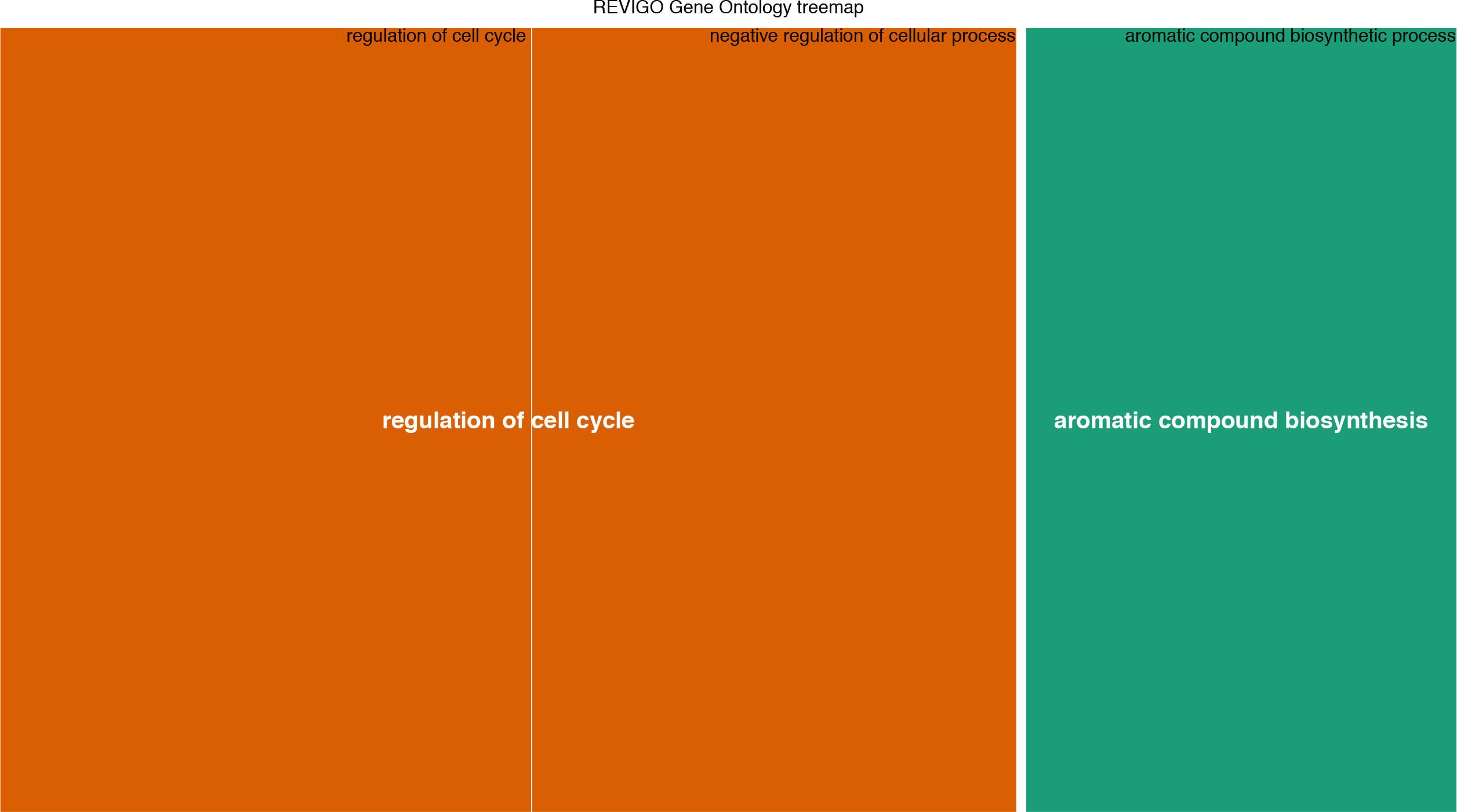
E. Coli cluster 3.

**SM Figure 8.**
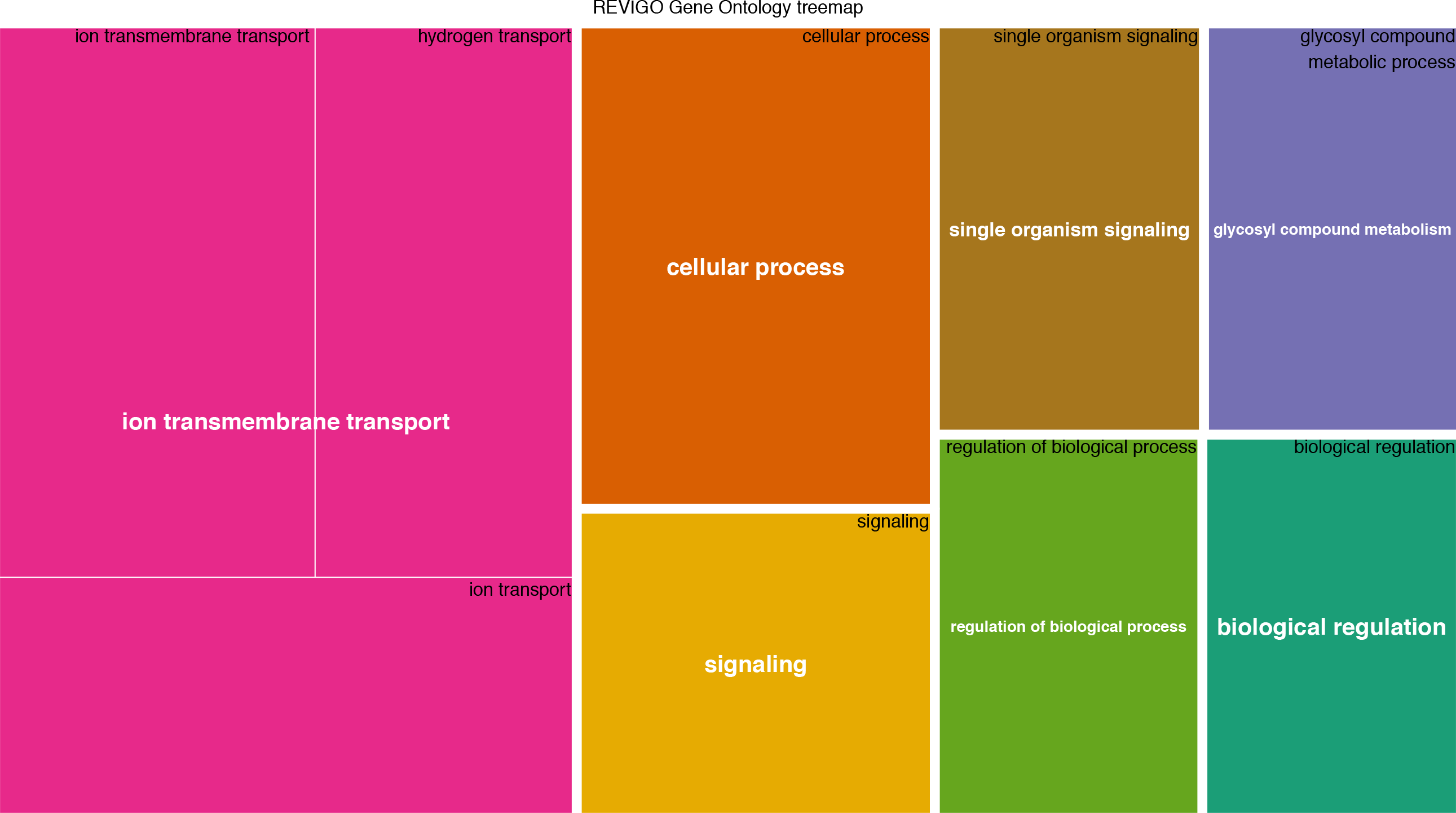
E. coli cluster 4.

**SM Figure 9.**
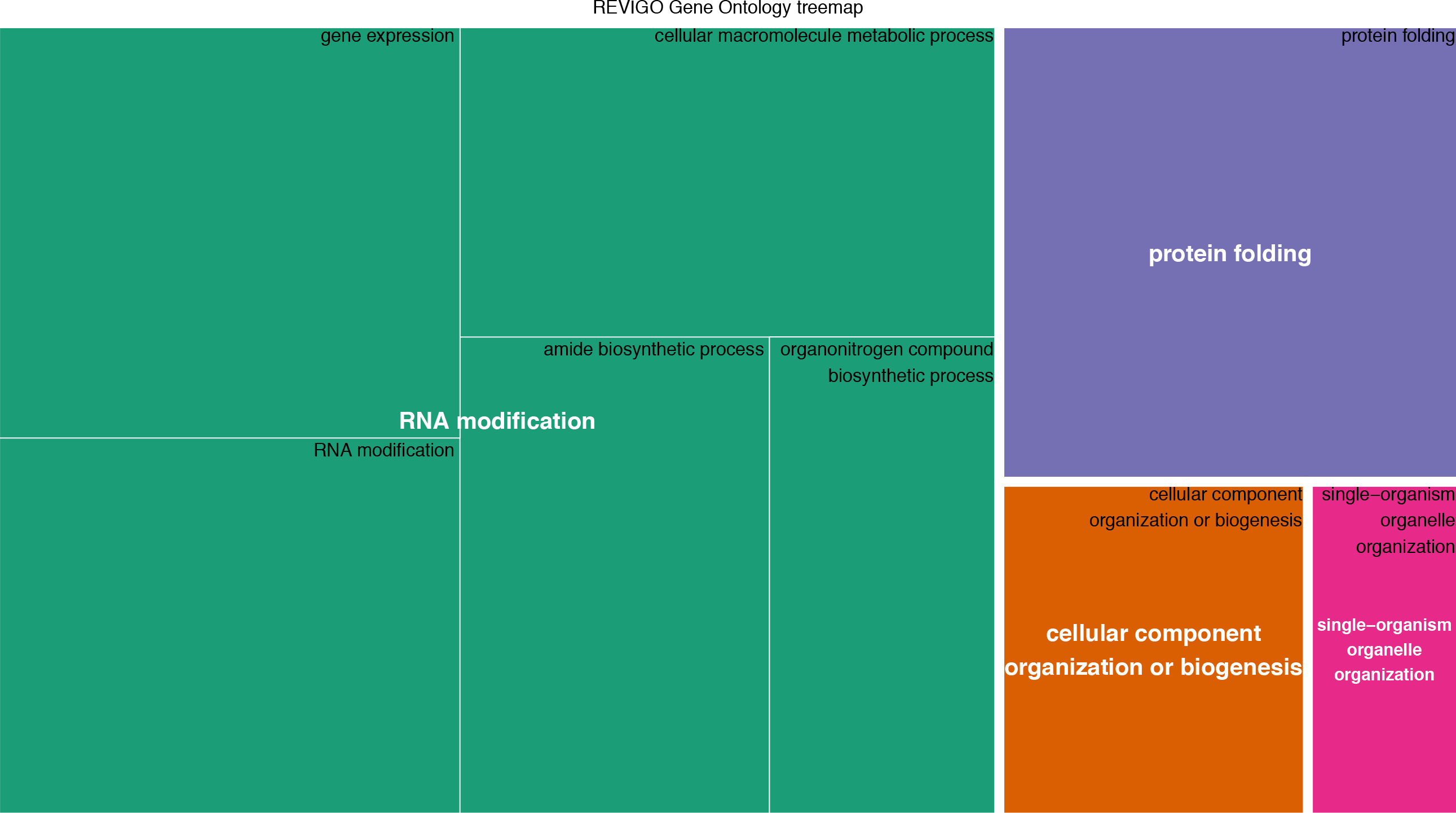
E. coli cluster 5.

**SM Figure 10.**
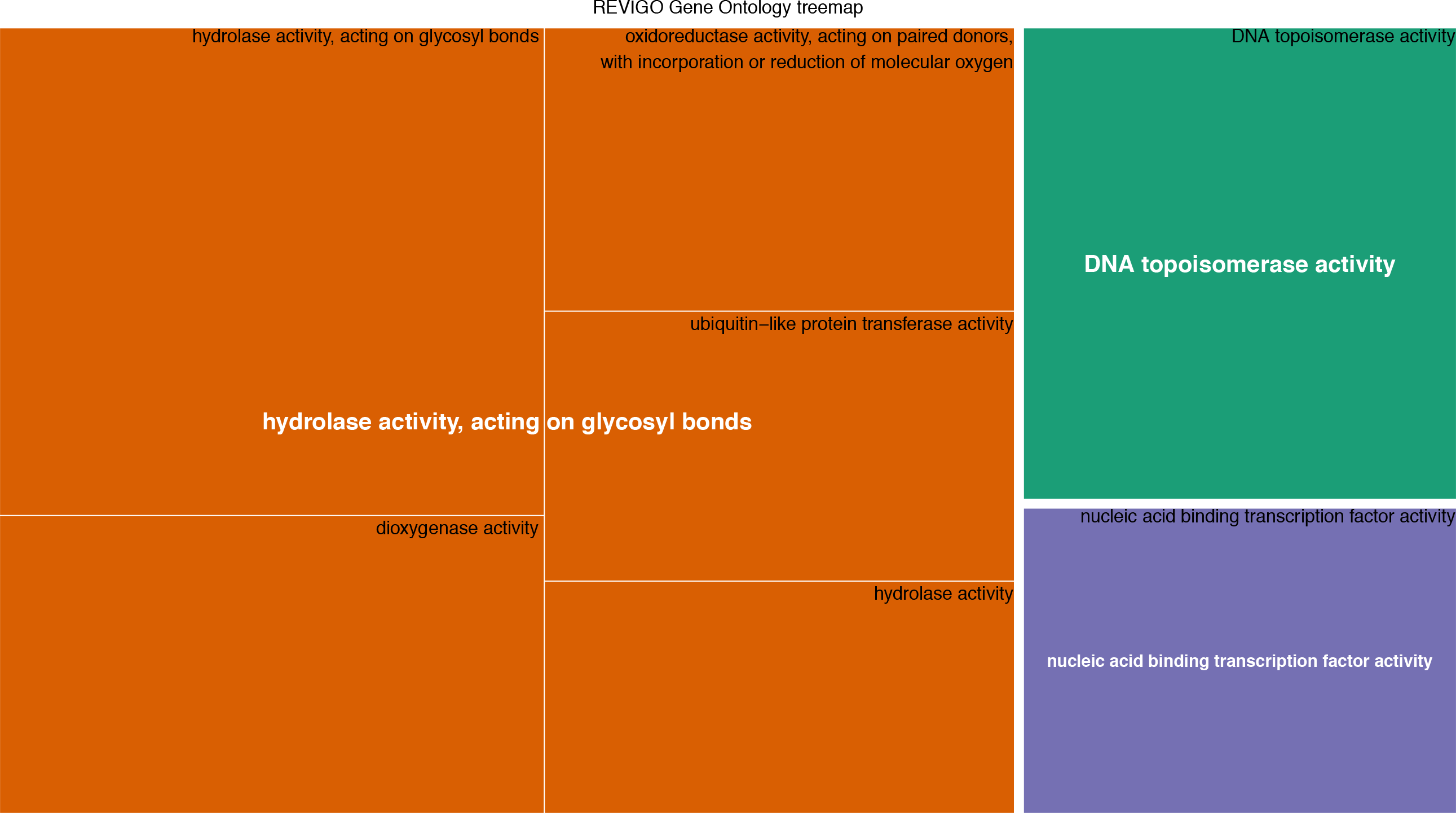
*E. coli* cluster 6.

**SM Figure 11.**
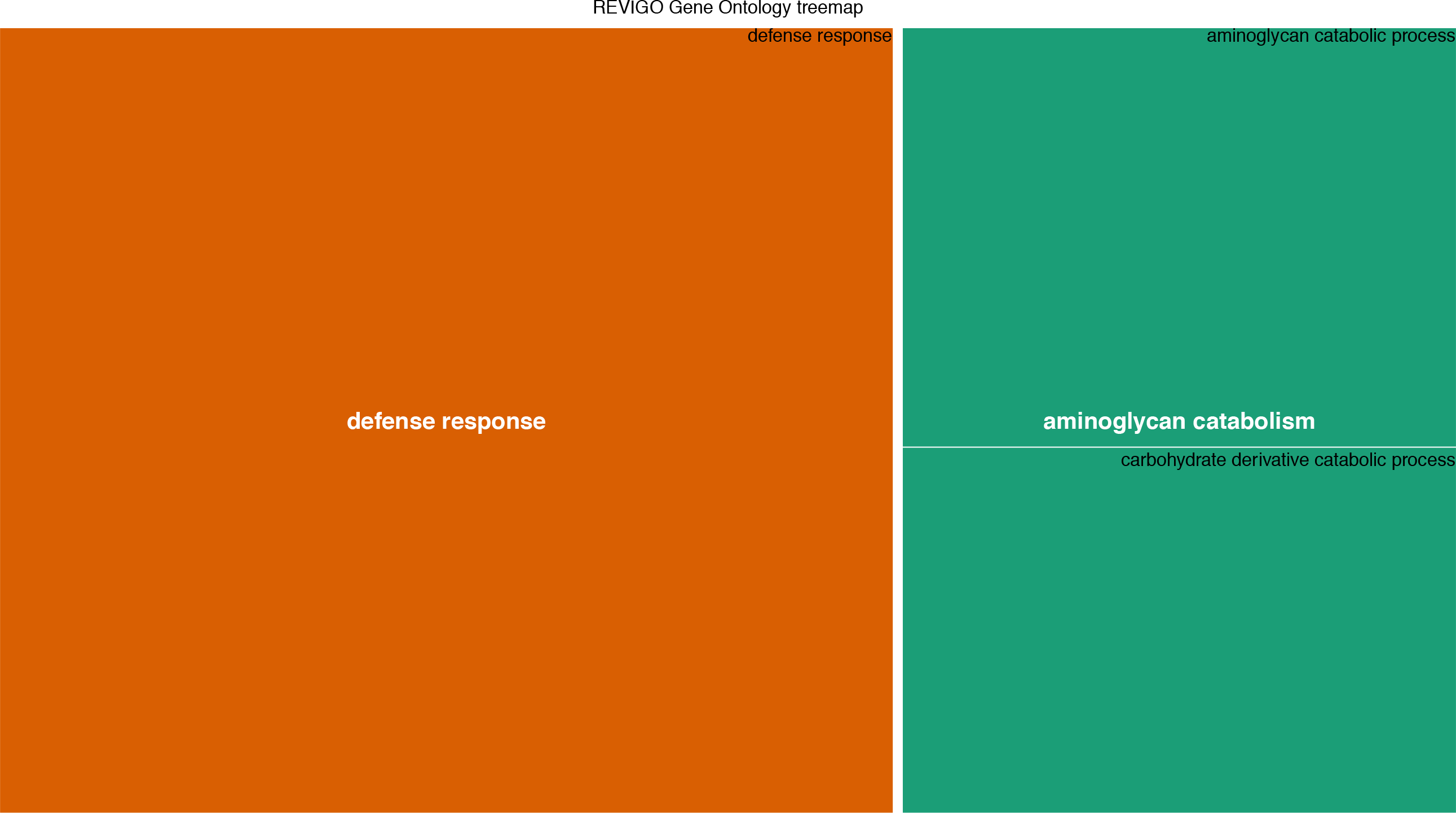
M. luteus cluster 1.

**SM Figure 12.**
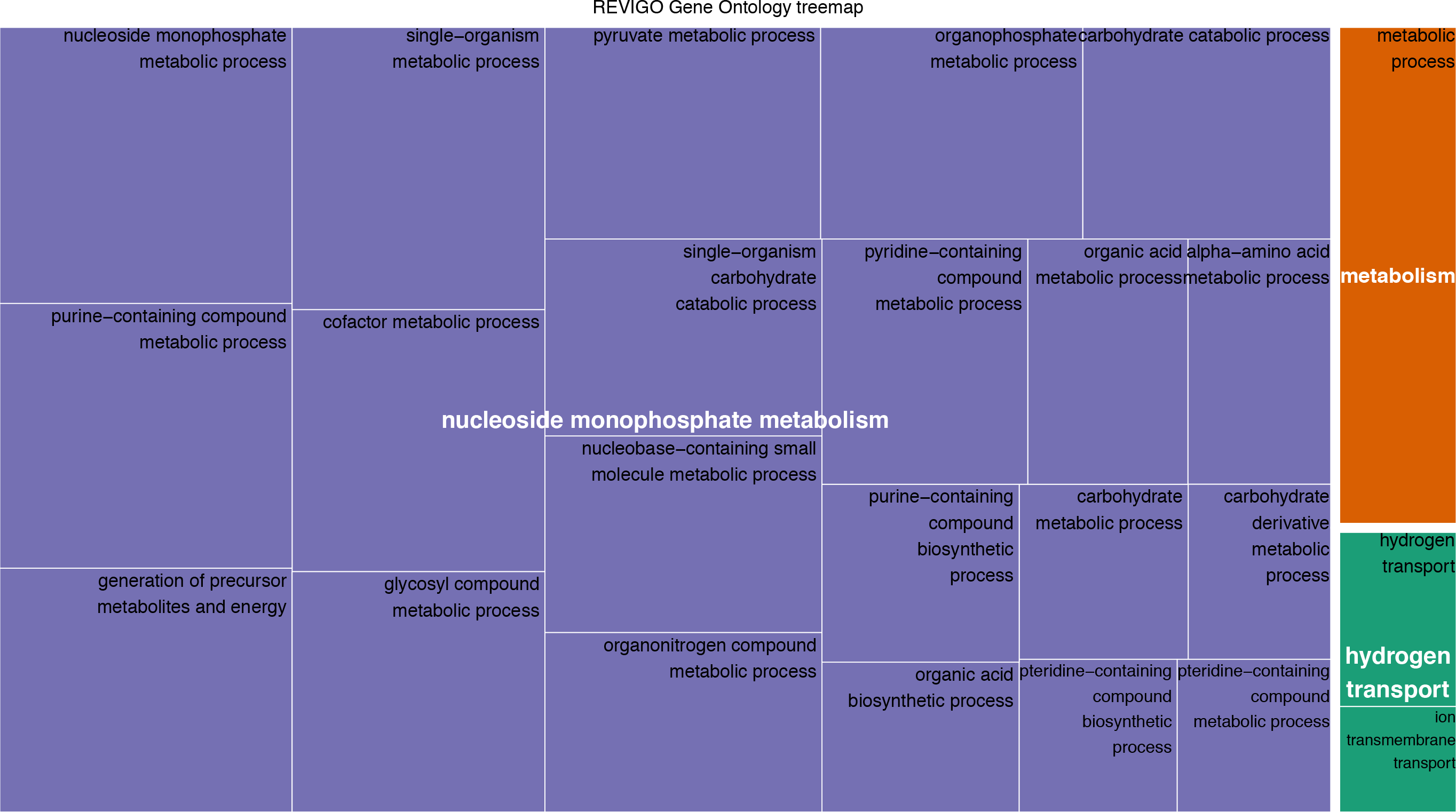
M. luteus cluster 2.

